# Branched-chain amino acid catabolism depends on GRXS15 through mitochondrial lipoyl cofactor homeostasis

**DOI:** 10.1101/2020.02.13.947697

**Authors:** Anna Moseler, Inga Kruse, Andrew E. Maclean, Luca Pedroletti, Stephan Wagner, Regina Wehler, Katrin Fischer-Schrader, Gernot Poschet, Markus Wirtz, Peter Dörmann, Tatjana M. Hildebrandt, Rüdiger Hell, Markus Schwarzländer, Janneke Balk, Andreas J. Meyer

## Abstract

Iron-sulfur (Fe-S) clusters are ubiquitous cofactors in all life and are used in a wide array of diverse biological processes, including electron transfer chains and several metabolic pathways. Biosynthesis machineries for Fe-S clusters exist in plastids, the cytosol and mitochondria. A single monothiol glutaredoxin (GRX) has been shown to be involved in Fe-S cluster assembly in mitochondria of yeast and mammals. In plants, the role of the mitochondrial homologue GRXS15 has only partially been characterized. Arabidopsis *grxs15* null mutants are not viable, but mutants complemented with the variant *GRXS15 K83A* develop with a dwarf phenotype. In an in-depth metabolic analysis, we show that most Fe-S cluster-dependent processes are not affected, including biotin biosynthesis, molybdenum cofactor biosynthesis and the electron transport chain. Instead, we observed an increase in most TCA cycle intermediates and amino acids, especially pyruvate, 2-oxoglutarate, glycine and branched-chain amino acids (BCAAs). The most pronounced accumulation occurred in branched-chain α-keto acids (BCKAs), the first degradation products resulting from deamination of BCAAs. In wild-type plants, pyruvate, 2-oxoglutarate, glycine and BCKAs are all metabolized through decarboxylation by four mitochondrial lipoyl cofactor-dependent dehydrogenase complexes. Because these enzyme complexes are very abundant and the biosynthesis of the lipoyl cofactor depends on continuous Fe-S cluster supply to lipoyl synthase, this could explain why lipoyl cofactor-dependent processes are most sensitive to restricted Fe-S supply in *GRXS15 K83A* mutants.

**One-sentence summary:** Deficiency in GRXS15 restricts protein lipoylation and causes metabolic defects in lipoyl cofactor-dependent dehydrogenase complexes, with branched-chain amino acid catabolism as dominant bottleneck.

## Introduction

Since the early days of biological evolution iron-sulfur (Fe-S) clusters have been employed as catalytic co-factors for electron transfer reactions and are nowadays present in a plethora of essential proteins (Pain and Dancis, 2016). Because Fe-S clusters are inherently instable they do not exist in free form but always need to be chaperoned before reaching their final destination apoproteins. Among the proteins thought to be involved in Fe-S cluster transfer downstream of the assembly machinery is a specific subtype of glutaredoxins (GRXs) capable of coordinating [2Fe-2S] clusters as a protein dimer (Banci et al., 2014; Couturier et al., 2015; Lill and Freibert, 2020).

Glutaredoxins are ubiquitous proteins, which form a large family with several subfamilies in plants (Rouhier et al., 2008; Meyer et al., 2009). Although their canonical function is glutathione-dependent redox catalysis, dissection of the function of subclasses and individual family members reveals an unexpectedly diverse picture (Lillig et al., 2008; Deponte, 2013). Class II GRXs share a CGFS amino acid motif in the active site and are proposed to serve as carrier proteins for Fe-S cluster between the assembly machinery and receiving apoproteins. A second proposed function is the repair of oxidation sensitive Fe-S clusters (Couturier et al., 2015). In Arabidopsis, Fe-S cluster assembly machineries are present in the cytosol, plastids and mitochondria and at least one monothiol GRX is located in each of these compartments: GRXS15 in mitochondria; GRXS14 and GRXS16 in plastids; and GRXS17 in the cytosol (Cheng et al., 2006; Bandyopadhyay et al., 2008; Moseler et al., 2015; Knuesting et al., 2015). While autonomous pathways for the multistep Fe-S protein maturation process are present in plastids and mitochondria, the cytosolic machinery relies on the export of bound sulfide as a precursor from mitochondria (Schaedler et al., 2014). While plants deficient in plastidic GRXS14 did not display any growth phenotype under non-stress conditions, genetic stacking of a *grxs14* null mutant and knockdown of GRXS16 caused pronounced growth retardation (Rey et al., 2017). Exposure of *grxs14* and the double mutant to prolonged darkness led to accelerated chlorophyll loss compared to wild type (WT) and decreased abundance of proteins involved in the maturation of Fe-S proteins. Mutants lacking the cytosolic GRXS17 were sensitive to high temperature and long-day photoperiod (Cheng et al., 2011; Knuesting et al., 2015). However, the activities of cytosolic Fe-S proteins, like aconitase (ACO) or aldehyde oxidase, were not substantially altered in *grxs17* null mutants (Knuesting et al., 2015; Iñigo et al., 2016).

The mitochondrial GRXS15 is indispensable as indicated by embryonic lethality of null mutants (Moseler et al., 2015). Partial complementation with a mutated *GRXS15 K83A* variant, which is weakened in its ability to coordinate an [2Fe-2S] cluster *in vitro*, results in a dwarf phenotype and diminished activity of the Fe-S protein ACO (Moseler et al., 2015). A similar dwarf phenotype has also been reported for a GRXS15 knockdown line, albeit without any effect on ACO activity (Ströher et al., 2016). Mitochondria contain at least 26 Fe-S proteins that are involved in different processes, including electron transport (complexes I, II and III in the respiratory electron transport chain) and the tricarboxylic acid (TCA) cycle [ACO and succinate dehydrogenase (SDH)]. A general role of GRXS15 in the early steps of Fe-S cluster transfer would therefore predict pleiotropic effects of diminished GRXS15 activity, due to the simultaneous impairment of several central mitochondrial processes. The number of potential defective sites is even further amplified if the synthesis of enzyme cofactors and the function of several cofactor-dependent enzymes, in turn, is compromised. Indeed, pathways for biosynthesis of the molybdenum cofactor (Moco) and lipoyl cofactor involve the mitochondrial [4Fe-4S] proteins GTP-3’,8-cyclase CNX2 (cofactor of nitrate reductase and xanthine dehydrogenase 2) and LIP1 (lipoyl synthase) (Yasuno and Wada, 2002; Schwarz and Mendel, 2006).

A pronounced decrease in lipoyl cofactor-dependent proteins in GRXS15 knockdown mutants led to the conclusion that efficient transfer of Fe-S clusters is required for mitochondrial lipoyl cofactor synthesis (Ströher et al., 2016). In the mitochondrial matrix, four enzyme complexes depend on lipoamide as a prosthetic group: the pyruvate dehydrogenase complex (PDC), the 2-oxoglutarate dehydrogenase complex (OGDC), the glycine decarboxylase complex (GDC), and the branched-chain α-keto acid dehydrogenase complex (BCKDC) (Taylor et al., 2004; Solmonson and DeBerardinis, 2018). The PDC acts as the entry point of acetyl-CoA into the TCA cycle, while OGDC acts within the TCA cycle to convert 2-oxoglutarate to succinyl-CoA. The GDC catalyzing the oxidative decarboxylation of glycine is essential for photorespiration (Douce et al., 2001), but also for C1 metabolism (Mouillon et al., 1999). BCKDC is involved in catabolism of the three branched-chain amino acids (BCAAs) leucine (Leu), valine (Val) and isoleucine (Ile) and their corresponding branched-chain α-keto acids (BCKAs) (Gu et al., 2010; Araújo et al., 2010; Peng et al., 2015). Whether all these lipoyl cofactor-dependent enzymes are affected similarly in *grxs15* mutants and whether other pathways containing Fe-S enzymes are diminished and thus constitute bottlenecks that severely restrict metabolic fluxes is yet unknown because the respective mutants have not been metabolically characterized.

Here, we aimed to identify the most severe metabolic bottlenecks caused by severely restricted capacity of GRXS15 mutants in Fe-S transfer. We consider several candidate Fe-S proteins involved in essential mitochondrial processes starting with biotin biosynthesis, followed by Moco biosynthesis, capacity of the mitochondrial electron transport chain, TCA cycle flow and closing with the biosynthesis of lipoyl cofactor. We assess how these Fe-S related processes are affected in *grxs15-3* null mutants complemented with *GRXS15 K83A* and in *GRXS15^amiR^* knockdown mutants trying to pin down the cause of the phenotype and by that the functional significance of GRXS15. By direct comparison of partially complemented null mutants and knockdown mutants we resolve previous contradictions about the role of GRXS15 in the maturation of Fe-S containing enzymes.

## Results

### *GRXS15 K83A* causes retardation in growth

To complete embryogenesis, GRXS15 is essential in plants. To bypass embryo lethality, Arabidopsis *grxs15* null mutants were complemented with the *GRXS15 K83A* variant which are able to grow, but the plants have small rosette leaves (Moseler et al., 2015). Based on that observation we aimed to further analyze the growth phenotype and compare with published records of *grxs15* knockdown mutants. A dwarf phenotype of the *GRXS15 K83A* complementation lines #1 to #5 becomes apparent at the early seedling stage (Fig. 1A, B). Analysis of root length in five randomly selected lines consistently also showed a concomitant reduction of primary root length compared to WT (Figure 1B).

**Figure 1.**
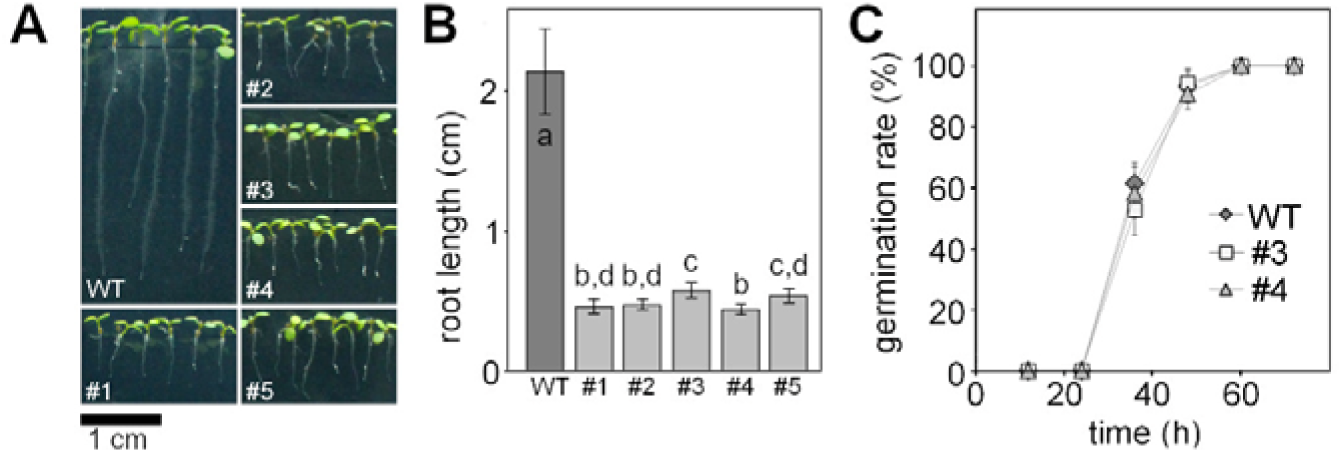
Complementation of the Arabidopsis *grxs15-3* mutant with *UBQ10_pro_:GRXS15 K83A*. **A:** 8-d-old wild-type (WT) seedlings compared with *GRXS15 K83A* mutants grown on vertical agar plates under long-day conditions. **B:** Primary root length of 8-d-old *GRXS15 K83A* mutants compared to WT (*n* = 35; means ± SD). Different letters indicate significant differences between the different lines; *P* ≤ 0.05; (one-way ANOVA with post hoc Holm-Sidak). **C:** Germination rate of *GRXS15 K83A* lines #3 and #4 compared to WT. All seeds were initially stratified at 4°C in the dark for 1 d (*n* = 6 with 20-25 seeds each; means ± SD). Germination was assessed with the emergence of the radicle. No statistically significant differences were found using Student’s t-Test analysis.

Although only minor differences in seedling size could be observed, line #3 was the best growing complementation line and line #4 the weakest (Fig.1C; Moseler et al., 2015). This effect was stable and consistent over several generations. The phenotype is similar to GRXS15amiR knockdown lines reported by Ströher et al. (2016) (Supplemental Fig. S1). A T-DNA insertion line grxs15-1 carrying a T-DNA in an intron within the 5’-UTR (Moseler et al., 2015), which had been reported to display a short root phenotype (Ströher et al., 2016) cannot be clearly distinguished from the WT in our hands, neither at seedling stage nor at rosette stage (Supplemental Fig. S1). This allele was excluded from further analysis. To test whether the reduced growth of GRXS15 K83A-complemented null mutants was true growth retardation or caused by delayed germination, the two lines #3 and #4 were scored for the timing of radical emergence. The absence of any difference between WT and the two mutants suggests that the growth phenotype reflects a genuine growth retardation (Fig. 1C).

### Biotin-mediated metabolism is not impaired when GRXS15 function is diminished

Following our earlier observation that GRXS15 can coordinate a [2Fe-2S] cluster (Moseler et al., 2015), similar to the closest homologs in yeast and mammals (Uzarska et al., 2013; Banci et al., 2014), we embarked on testing a number of pathways of Fe-S-dependent metabolism that may be affected in the mutant. One putative target protein of GRXS15 is mitochondrial biotin synthase (BIO2, At2g43360) since it relies on supply of a [2Fe-2S] and a [4Fe-4S] cluster. BIO2 catalyzes the final step in biotin biosynthesis, which acts as an essential cofactor in several carboxylases in energy metabolism. Destruction of the [2Fe–2S] cluster for sulfur supply to biotin with each catalytic cycle and subsequent turnover increases the demand for [2Fe-2S] clusters (Ugulava et al., 2001). *bio2* null mutants were previously described as embryo-defective, arrested mostly at globular or heart stage of embryo development (Patton et al., 1998; Meinke, 2019). Because lack of biotin typically causes degradation of the respective apoproteins (Solbiati et al., 2002), we tested for the abundance of biotin-dependent methylcrotonoyl-CoA carboxylase (MCCase), which is involved in leucine degradation in mitochondria. None of the five analyzed *grxs15* complementation lines showed a decrease in protein abundance of the biotinylated MCCase subunit A (MCCA) (Fig. 2A). Biotin is also exported to the cytosol and the chloroplasts, where it is required for synthesis and elongation of fatty acids by hetero- and homomeric acetyl-CoA carboxylase (ACCase). Total fatty acids in seeds amounted to 7.6 ± 0.8 nmol seed^-1^ in line #4 and 7.6 ± 1.0 nmol seed^-1^ in the WT and no difference in relative abundance of specific fatty acids in seeds was observed (Fig. 2C). In 8-day-old seedlings the amount of total fatty acids did not differ in line #4 10.3 ± 0.4 nmol (mg FW)^-1^ compared to 8.8 ± 1.0 nmol (mg FW)^-1^ in WT, but a 23% increase in α-linolenic acid (18:3) was observed (Fig. 2B).

**Figure 2.**
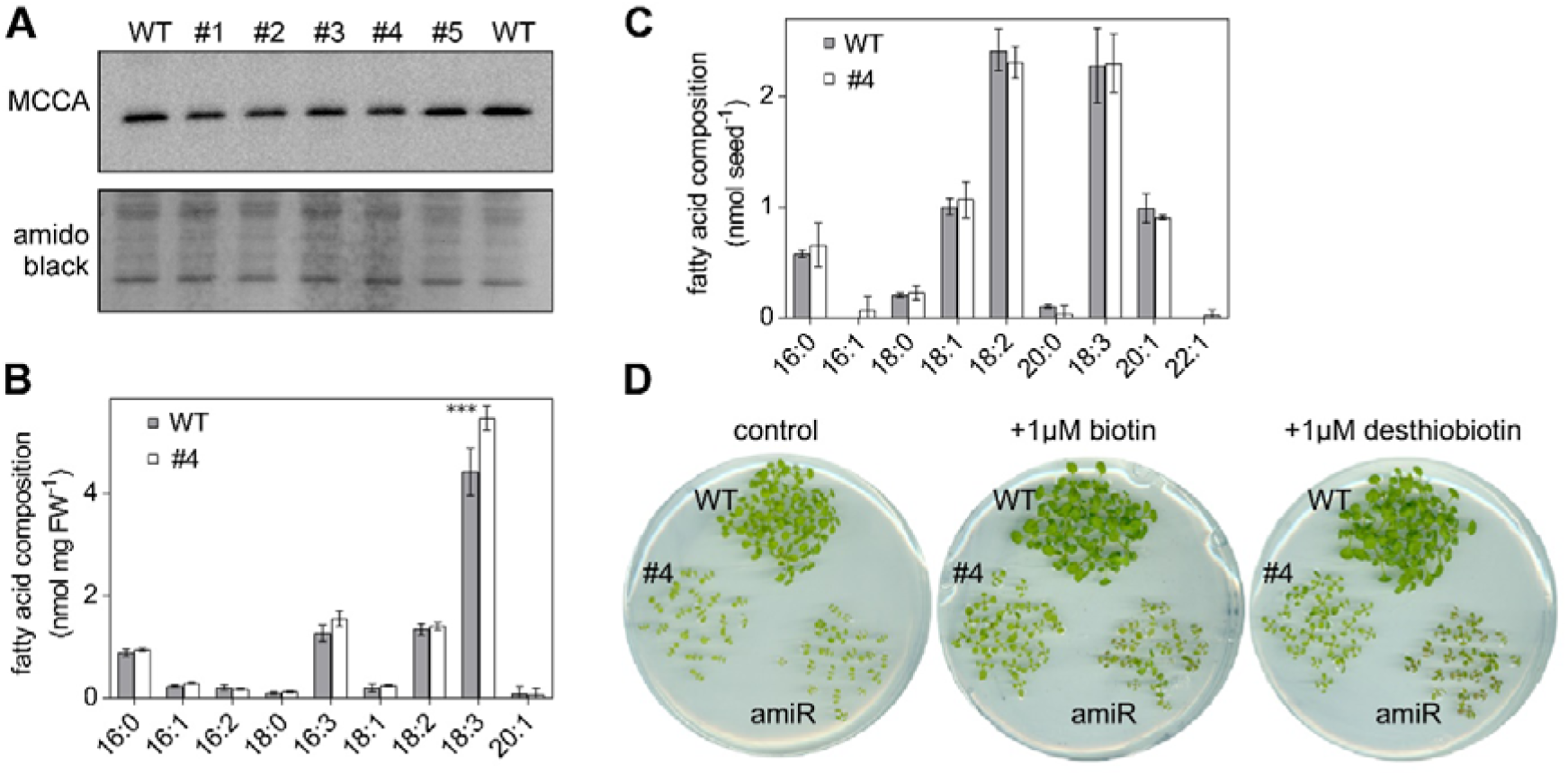
*GRXS15 K83A* mutation has no impact on the biotin pathway in Arabidopsis seedlings. **A:** Immunoblot analysis of biotinylated MCCA in mitochondria of *GRXS15 K83A* mutants compared with WT. In the upper panel, biotinylated MCCA was detected by streptavidin HRP in isolated mitochondria from 2-weeks-old seedlings (9 μg protein was loaded per lane). In the lower panel, amido black staining of the membrane is shown as a control for protein loading. **B, C:** Fatty acids quantified by gas chromatography using a flame ionization detector of 8-d-old seedlings (B) and seeds (C) of *GRXS15 K83A* line #4 compared to WT (*n* = 3-4; means ± SD). The statistical analysis (two-way ANOVA with post hoc Holm-Sidak comparisons for WT vs. *grxs15*) indicated no significant (*P* ≤ 0.05) change except for 18:3 (*** = P < 0.001). **D:** *GRXS15 K83A* line #4, the knockdown line *GRXS15^amiR^* (amiR) and wild-type plants were grown on horizontal plates with ½ MS agar without sucrose. The medium contained either no biotin (control), 1 μM biotin or 1 μM desthiobiotin.

*bio2* mutants can be rescued by the addition of biotin to both arrested embryos cultured *in vitro* and to mutant plants grown on soil (Schneider et al., 1989; Patton et al., 1998; Pommerrenig et al., 2013). External supply of biotin or its precursor desthiobiotin to a *GRXS15^amiR^* knockdown mutant and the complemented line #4 in both cases improved growth slightly but did not rescue the growth defects of either of the lines (Fig. 2D). It should be noted though that also the WT grew better with supply of biotin or desthiobiotin. These results suggest that growth retardation of *grxs15* mutants is not primarily caused by defects in biotin synthesis.

### Moco-dependent nitrogen metabolism is not limiting upon impaired GRXS15 function

The Moco precursor cyclic pyranopterin monophosphate (cPMP) is synthesized in the mitochondrial matrix by CNX2 (At2g31955) and the cyclic pyranopterin monophosphate synthase CNX3 (At1g01290) and is exported to the cytosol for subsequent biosynthesis steps (Bittner, 2014; Kruse et al., 2018). Because CNX2 contains two [4Fe-4S] clusters, we hypothesized that Moco biosynthesis and hence Moco-dependent biochemical pathways may be affected by defects in mitochondrial Fe-S transfer. The most abundant Moco-dependent enzymes include nitrate reductase (NR), aldehyde oxidase (AO), xanthine dehydrogenase (XDH) and sulfite oxidase (SO). Arabidopsis generally prefers nitrate as nitrogen source (Sarasketa et al., 2014), but mutants deficient in Moco biosynthesis can be rescued by providing ammonium as a nitrogen source to bypass nitrate reductase (Wang et al., 2004; Kruse et al., 2018), revealing NR as the main recipient of Moco. While the preference for nitrate (KNO_3_) over ammonium ((NH_4_)_2_SO_4_) could be confirmed in wild-type plants, we found that the growth retardation of *GRXS15 K83A* roots is more pronounced on nitrate than on ammonium as sole nitrogen source (Fig. 3A). Similar results were obtained when seedlings were grown on NH_4_Cl instead of (NH_4_)_2_SO_4_ to control for possible impacts of the respective counter anions on the growth behavior (Supplemental Fig. S2A).

**Figure 3.**
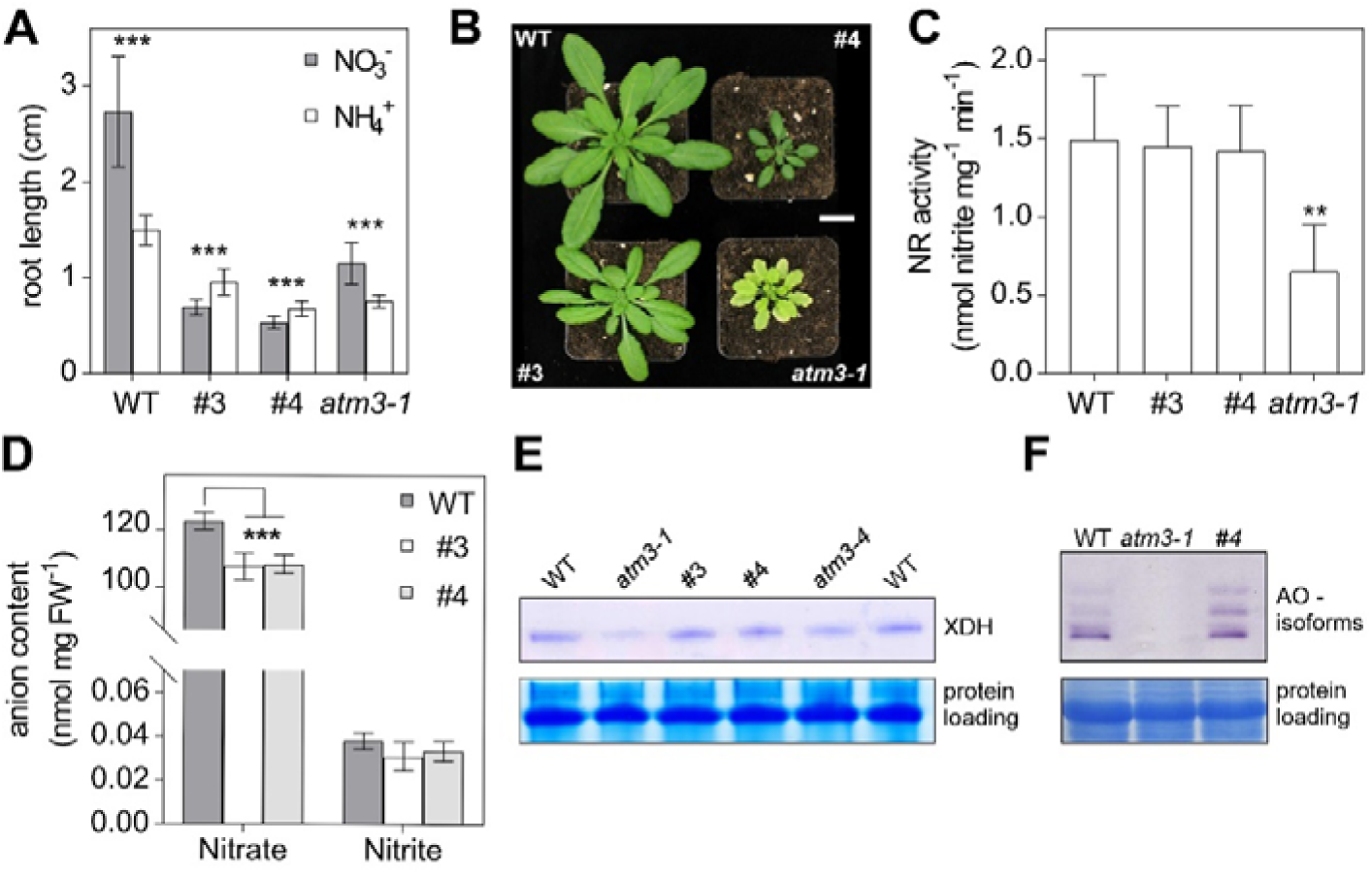
Growth of Arabidopsis *GRXS15 K83A* mutants is affected by the nitrogen source. **A:** Primary root length of *GRXS15 K83A* lines #3 and #4 as well as *atm3-1* seedlings compared to WT grown on vertical agar plates containing 5 mM KNO_3_ or 2.5 mM (NH_4_)_2_SO_4_ as N-source for 8 d under long-day conditions (*n* = 30; means ± SD). Student’s t-Test analysis showed significant differences between the growth on the different inorganic N-sources in all lines ***: *P* < 0.001. **B:** Representative 4-week-old plants of WT, *GRXS15 K83A* lines #3 and #4 and *atm3-1* all grown on soil under long-day conditions. Scale bar = 2 cm. **C:** Nitrate reductase activity in WT, lines #3 and #4 as well as in *atm3-1*. Activity was analyzed in 4-week-old plants grown on soil by measuring the presence of nitrite via the Griess reaction (*n* = 4; means ± SD, **: *P* ≤ 0.01). **D:** Nitrate and nitrite content of 8-d-old WT and *GRXS15 K83A* lines #3 and #4 seedlings grown on agar plates (*n* = 4; means ± SEM). The statistical analysis (two-way ANOVA with post hoc Holm-Sidak comparisons for WT vs. *grxs15*) indicated a significant change in the nitrate content; ***: *P* ≤ 0.001. **E:** In-gel activity of XDH in WT, *atm3-1,* and *GRXS15 K83A* mutants. Equal amounts of protein (35 μg) extracted from 8-d-old seedlings were separated on non-denaturing PA gel and stained for XDH activity using hypoxanthine as substrate. **F:** In-gel activities of aldehyde oxidase (AO) in WT and *atm3-1* as well as *grxs15* mutants. Equal amounts of protein were separated on non-denaturing PA gels and stained for AO activity using synthetic aldehydes (1-naphthaldehyde and indole-3-carboxaldehyde) as substrates. For control of protein-loading the gel was subsequently stained with Coomassie.

The pronounced growth retardation on nitrate could be indicative of severe NR deficiency similar to *nia1 nia2* mutants lacking functional NR (Wilkinson and Crawford, 1993). A similar NR deficiency has been described for mutant alleles of the ABC transporter ATM3 that is involved in Moco biosynthesis (Bernard et al., 2009; Teschner et al., 2010; Kruse et al., 2018). *atm3-1* mutants display a severe growth phenotype and are chlorotic (Fig. 3B). While *GRXS15 K83A* mutants are also smaller than WT, they are not chlorotic and thus do not phenocopy *atm3-1* (Fig. 3A, B). Despite NR activity being diminished to 50% of WT, root growth of *atm3-1* was still better on nitrate than on ammonium (Fig. 3A, C). NR activity was not altered in the *GRXS15 K83A* mutants #3 and #4 (Fig. 3C). Despite the unaffected NR activity, both *grxs15* mutants contained significantly less nitrate than WT seedlings (Fig. 3F). Nitrite and other inorganic anions like chloride, sulfate or phosphate were not altered between the mutant lines and WT (Supplemental Fig. S2B). All other tested Moco-dependent enzymes such as AO or XDH showed no decrease in activity in the *grxs15* mutants compared to WT (Fig. 3E, F). Taken together, these results suggest that NR activity in *GRXS15 K83A* mutants is sufficient to use nitrate as the sole nitrogen source and does not explain the growth inhibition on nitrate.

### Impaired GRXS15 function leads to decreased root respiration

The mitochondrial *electron* transport chain (mETC) contains three enzyme complexes with a total of 12 Fe-S cofactors: complex I with two [2Fe-2S] and six [4Fe-4S] clusters, complex II with one [2Fe-2S], one [3Fe-4S], and one [4Fe-4S] cluster, and complex III with one [2Fe-2S] cluster (Couturier et al., 2015; Meyer et al., 2019). Thus, we measured the respiration of detached roots and dissected the capacity of complex I and II-linked electron flow. Indeed, roots of line #3 displayed a decreased respiration rate of 1.31 ± 0.35 nmol O_2_ min^-1^ (mg DW)^-1^ compared with the wild-type rate of 2.92 ± 0.62 nmol O_2_ min^-1^ (mg DW)^-1^ (Fig. 4A). This is similar to root tips of *GRXS15^amiR^* knockdown plants which were reported to consume less oxygen than wild-type plants (Ströher et al., 2016). Addition of the cytochrome *c* oxidase inhibitor KCN decreased the rate of both lines down to similar values. The remaining rates are accounted for by the presence of alternative oxidases (AOXs), since they could be inhibited by propylgallate (pGal). Interestingly, the AOX capacity appeared unchanged in line #3, even though AOX is highly inducible by mitochondrial dysfunction. Next, we investigated if the decreased root respiration is due to defects in the respiratory machinery or due to restricted metabolite turnover, or both. First, we compared the abundance of respiratory complexes in isolated mitochondria from *GRXS15 K83A* line #4, *GRXS15^amiR^* by BN-PAGE. None of the respiratory complexes including the Fe-S cluster containing complexes I, II and III was decreased in abundance in either mutant (Fig. 4B). Additionally, we purified mitochondria from whole seedlings of the *GRXS15 K83A* line #3 and supplemented them with succinate or pyruvate/malate, respectively, as respiratory substrates. Succinate provides electrons to the ubiquinone pool of the mETC via complex II, whereas pyruvate/malate predominantly provides NAD(P)H mainly generated by malate dehydrogenase and the PDC. NADH is subsequently oxidized mainly by complex I of the mETC and NAD(P)H by matrix-exposed alternative NADH-dehydrogenases. No differences in the respiration of isolated mitochondria were found with supply of succinate or pyruvate/malate (Fig. 4C, D), suggesting that the differences in respiration observed in whole roots cannot be accounted for by decreased capacities of the Fe-S cluster-containing complexes. In summary, similar total respiratory activities of WT and mutants further indicate that the *in vivo* difference in respiration rate is not due to a defect at the level of the mETC, but rather upstream or downstream.

**Figure 4.**
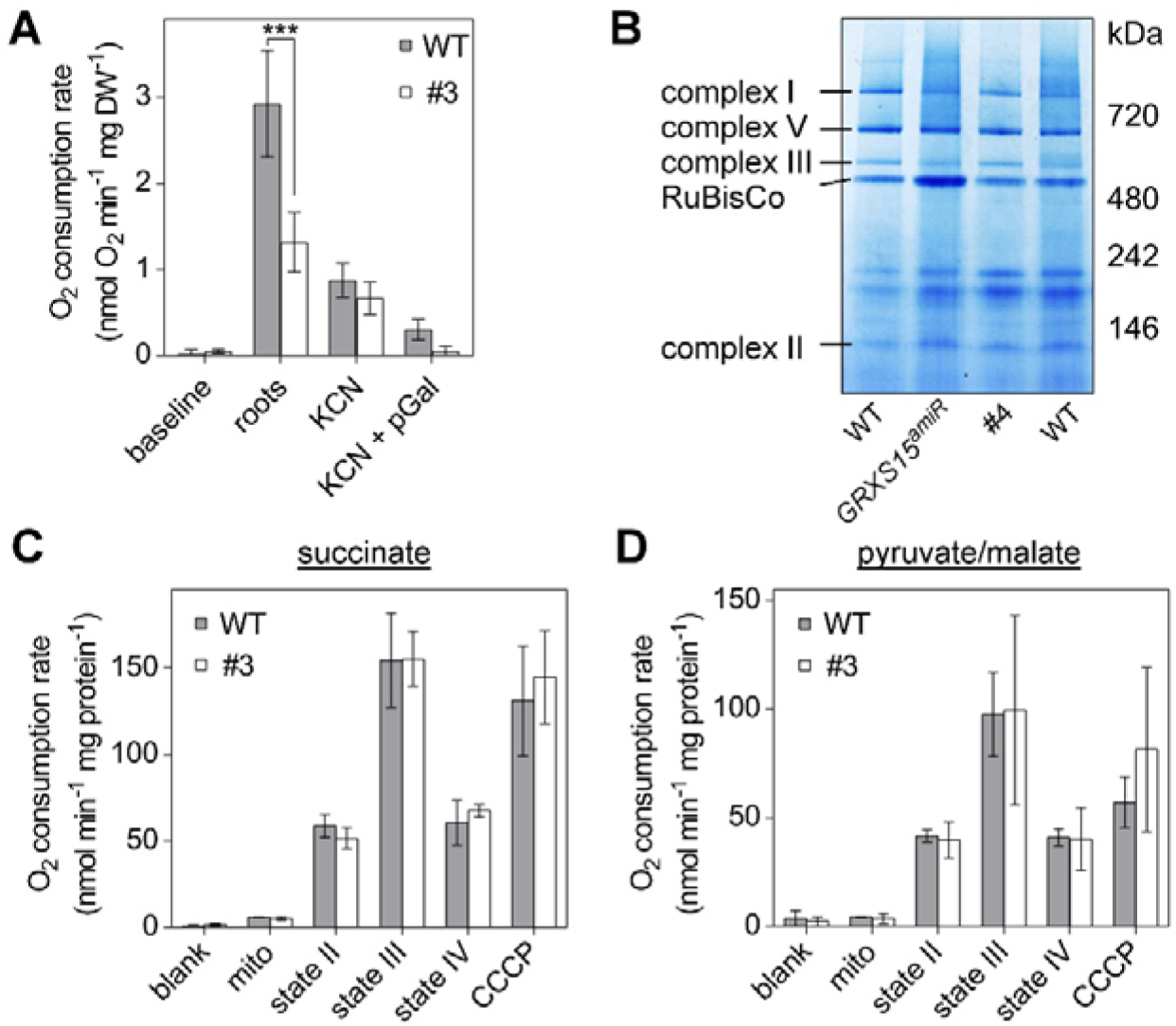
Respiration in complemented Arabidopsis *grxs15* mutants. **A:** Root respiration rate of *GRXS15 K83A* line #3 (4.5-week-old) and the respective WT grown to similar size (2-week-old) after addition of the cytochrome c oxidase inhibitor KCN (4 mM) alone or together with the alternative oxidase inhibitor propylgallate (pGal; 0.2 mM) (*n* = 4; means ± SD). The statistical analysis (two-way ANOVA with post hoc Holm-Sidak comparisons for WT vs. *grxs15* mutant) indicated a significant difference in the respiration of mitochondria from WT and *GRXS15 K83A* line #3; ***: *P* ≤ 0.001. **B:** Respiratory complexes I, II, III and V separated by BN-PAGE and visualized with Coomassie staining in WT, *GRXS15 K83A* line #4 and *GRXS15^amiR^*. Mitochondria were purified from 4-week-old plants. **C, D:** Oxygen consumption rates for purified mitochondria from WT *and GRXS15 K83A* line #3 energized with succinate or pyruvate/malate. O_2_ consumption was measured before (blank) and after addition of mitochondria (mito). State II respiration was initiated by the addition of the respective substrate (state II; succinate (10 mM succinate, 0.25 mM ATP) or pyruvate/malate (10 mM pyruvate, 10 mM malate, 0.3 mM NAD and 0.1 mM thiamine pyrophosphate). State III respiration was initiated by the addition of 50 μM ADP. State IV represents the respiration after ADP consumption and CCCP shows the respiration after addition of the protonophore carbonyl cyanide m-chlorophenylhydrazone (CCCP; 10 μM), which uncouples electron transport from ATP synthesis. All results are based on three independent preparations of mitochondria and are shown as means ± SEM.

The *capacity* for electron flow in isolated mitochondria does not allow conclusions about the actual mETC *activity in planta*. Hence, we tested whether the decreased respiration rate may result in a change of the ATP status of the cells. For analyses of the MgATP^2-^ level wild-type plants as well as the *grxs15* mutants #3 and #4 were transformed with the MgATP^2-^ biosensor ATeam1.03-nD/nA (De Col et al., 2017) targeted to the cytosol. As cytosolic ATP is predominantly provided by the mitochondria (Igamberdiev et al., 2001; Voon et al., 2018), any disturbance in the mitochondrial ATP synthesis will also affect the ATP level in the cytosol. Similar to the report by De Col et al. (2017) higher Venus/CFP fluorescence ratios indicating more efficient FRET between the sensor subunits and hence higher MgATP^2-^ levels were found in cotyledons compared to roots (Supplemental Fig. S3). However, no differences in the Venus/CFP emission ratio could be observed between WT and *GRXS15 K83A* mutants indicating similar cytosolic ATP levels (Supplemental Fig. S3). It should be noted though that the energy charge of the adenylate pool cannot be deduced from these results as it would require also analysis of AMP and ADP.

Previously we reported a 60% decrease in aconitase activity (Moseler et al., 2015), which at last partially explain the decreased respiration rate, but a decrease in aconitase was not seen in Ströher *et al*., 2016. To clarify the situation, we measured the activity of ACO, a [4Fe-4S] enzyme, in the K83A and amiRNA mutants grown under the same conditions side by side. Despite similar amounts of ACO protein in mitochondria of WT and the mutants *GRXS15^amiR^* and *GRXS15 K83A #4*, ACO activity was decreased to approximately 40% in isolated mitochondria of both mutants (Fig. 5). The observation that ACO activity is decreased, while the ACO protein abundance is the same, is surprising because it is generally assumed that ACO apoproteins are rather unstable and would be degraded (Castro et al., 2019). Either, the ACO protein is stabilized in a yet unknown manner or ACO activity is compromised in another way.

**Figure 5.**
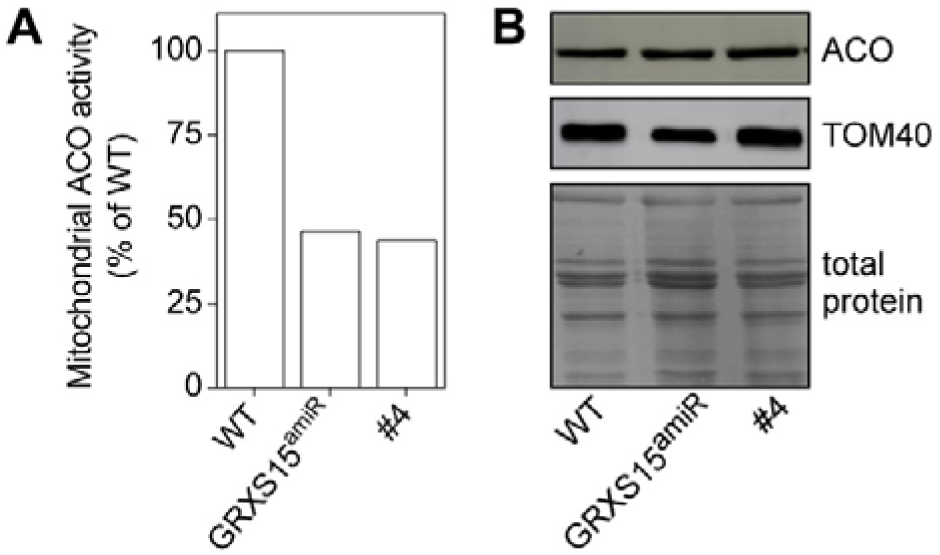
Aconitase activities in mitochondria of *grxs15* mutants. **A:** Aconitase activity of *GRXS15^amiR^* and G*RXS15 K83A* line #4 compared to the respective WTs from isolated mitochondria. *n* = 2. **B:** Protein gel blot analysis probed with antiserum raised against Arabidopsis ACO. 9 µg of protein isolated from mitochondria of a wild-type plant as well as *GRXS15^amiR^* and G*RXS15 K83A* lines #4 were loaded onto the gel. ACO and translocase of the mitochondria 40 (TOM40) protein levels were visualized by immunoblotting under denaturing conditions. Total protein staining served as a loading control.

### Diminished GRXS15 activity does not lead to any major signs of oxidative stress

Yeast *Δgrx5* mutant as well as a Arabidopsis *grxs14* null mutant are sensitive to oxidative stress and at least for the *Δgrx5* it was shown that specific proteins are oxidized in this mutant (Rodríguez-Manzaneque et al., 1999; Cheng et al., 2006). Aconitase is highly sensitive to oxidative stress and redox metabolism in the matrix (Verniquet et al., 1991; Navarre et al., 2000; Castro et al., 2019; Nietzel et al., 2020), suggesting that lower ACO activities may result from iron-mediated ROS formation as a possible consequence of an improper Fe-S cluster transfer by the *GRXS15 K83A* variant. However, staining of leaves with DAB for H_2_O_2_ and NBT for superoxide revealed no differences between WT and *grxs15* mutants (Supplemental Fig. S4). Since histological stains only provide a crude indication of major changes in ROS dynamics, but are not sufficiently sensitive to resolve localized intracellular changes in oxidant load, we next analyzed mitochondria-specific changes in H_2_O_2_ concentration or the glutathione redox potential (*E*_GSH_). The genetically encoded sensors roGFP2-Orp1 (Nietzel et al., 2019) and roGFP2-hGrx1 (Albrecht et al., 2014) were expressed in the mitochondrial matrix of both WT and mutant plants. Both sensors were highly reduced under control conditions and neither roGFP2-Orp1 nor roGFP2-hGrx1 revealed any significant differences between WT and *GRXS15 K83A* mutants in mitochondria of cotyledons and root tips (Supplemental Fig. S4B, C). Both roGFP2-sensor variants remained highly reduced in all lines as indicated by similar fluorescence ratios that resembled those after incubation with DTT for full sensor reduction. This indicates no major oxidative challenge in the mitochondrial matrix. Both sensors were responsive to oxidative challenge as indicated by a pronounced ratio change upon H_2_O_2_ addition.

### Diminished GRXS15 activity leads to accumulation of TCA cycle intermediates

To investigate any other metabolic defects in the GRXS15 K83A mutant, we measured the concentrations of several organic acids in the *GRXS15 K83A* mutants. We found each of the analyzed organic acids in the complemented *grxs15* mutants #3 and #4 to be increased. Pyruvate showed the most pronounced change, increasing by more than four-fold from 31.5 ± 2.4 pmol (mg FW)^-1^ in the WT to 131.76 ± 3.8 and 153.97 ± 16.5 pmol (mg FW)^-1^ in line #3 and #4 (Fig. 6). The accumulation of citrate and isocitrate was significant in line #4, but not in line #3. 2-oxoglutarate and malate showed minor increases in line #3 and pronounced increases in line #4. This trend did not reach statistical significance, however. A similarly concerted accumulation of TCA cycle intermediates was previously observed in antisense lines of the mitochondrial manganese superoxide dismutase 1 (MSD1) (Morgan et al., 2008). Those lines showed impaired mitochondrial ACO activity to less than 50%, suggesting that the compromised ACO activity is sufficient as an explanation for the rearrangements in the pools of TCA cycle intermediates. However, pyruvate content was not determined in the *MSD1 antisense* lines and the increased pyruvate content found in *GRXS15 K83A* lines cannot be straightforwardly linked to ACO activity.

**Figure 6.**
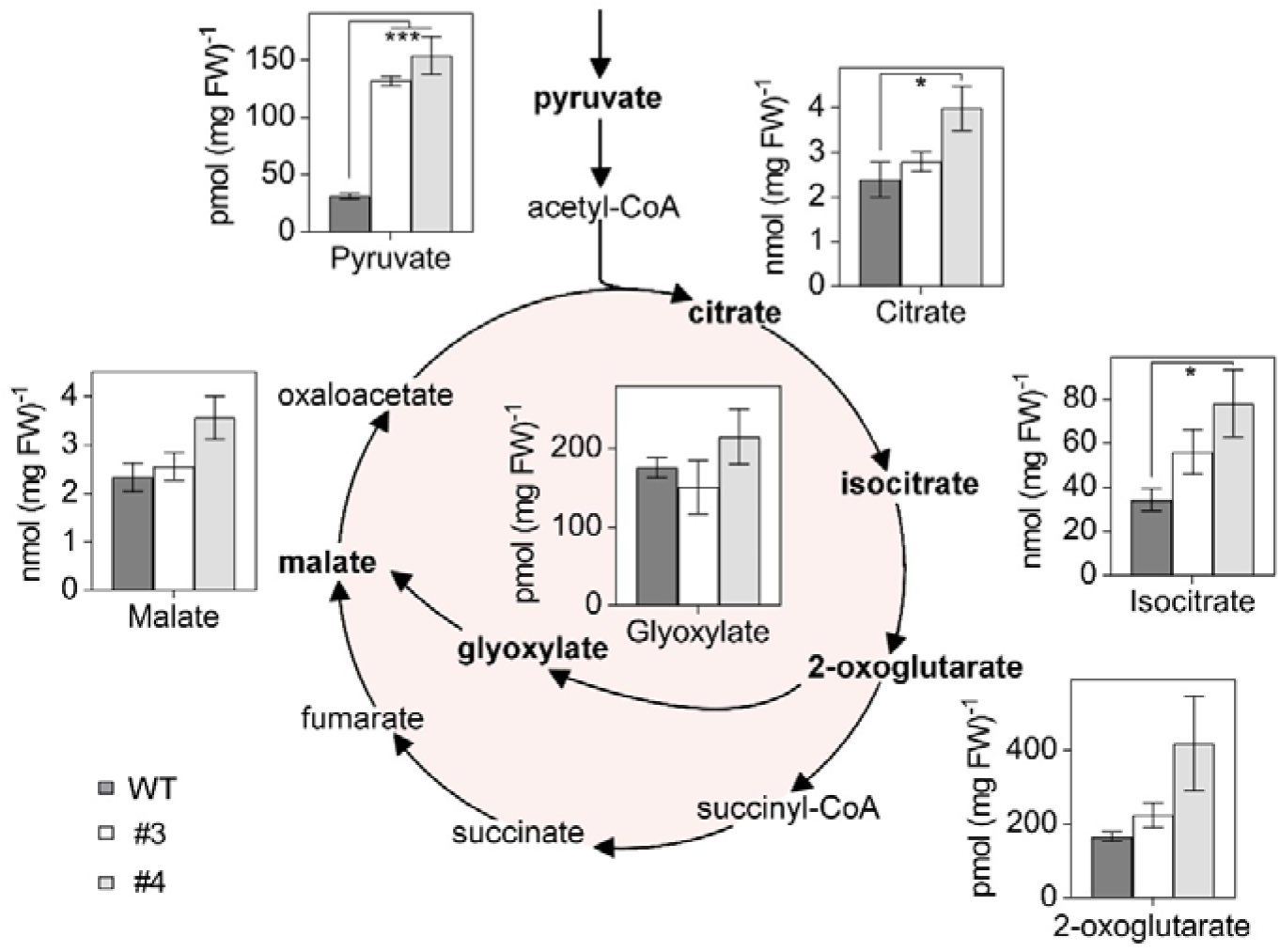
Organic acids of the TCA cycle accumulate in Arabidopsis *GRXS15 K83A* mutants. Organic acids were analyzed in 8-d-old seedlings of WT compared to *GRXS15 K83A* lines #3 and #4 (*n* = 4-5; means ± SEM). The statistical analysis (one-way ANOVA with post hoc Holm-Sidak comparisons for WT vs. mutant lines) indicated significant changes; *: *P* ≤ 0.05; ***: *P* ≤ 0.001.

### Alterations in pyruvate and glycine metabolism are correlated with impairment of lipoyl cofactor-dependent enzymes under diminished GRXS15 activity

The pronounced pyruvate accumulation may be caused by a backlog of metabolites due to a lower TCA flux or by a diminished activity of PDC, which catalyzes the decarboxylation of pyruvate to acetyl-CoA (Yu et al., 2012). The E2 subunit of this multi-enzyme complex uses a lipoyl cofactor, the synthesis of which was shown to be compromised in *GRXS15^amiR^* mutants (Ströher et al., 2016). In plant mitochondria, the lipoyl moiety is an essential cofactor of four protein complexes: PDC, OGDC, BCKDC, and GDC (Taylor et al., 2004). Ströher et al. (2016) showed decreased lipoylation of PDC E2-2 and E2-3 but no effects on E2-1. On the other hand, a pronounced decrease was observed in all GDC H protein isoforms and differences in the degree of lipoylation were explained by different modes of lipoylation. To get insight into the metabolic effects of diminished GRXS15 activity, we tested for protein lipoylation in the weakest complementation line #4 and directly compared the results to lipoylation in *GRXS15^amiR^* and WT. Furthermore, the complementation lines #3 and #4 were characterized for metabolites dependent on lipoyl cofactor-dependent enzymes. Immunodetection of the lipoyl group with specific antibodies to the cofactor indicated that the amount of lipoate bound to the H subunit isoforms of GDC was decreased in the *GRXS15 K83A* mutant to a similar extent as in *GRXS15^amiR^* (Fig. 7A). In contrast, the H protein levels were largely unchanged in all tested lines. GRXS15 was barely detectable in *GRXS15^amiR^* while in line #4 the mutated *GRXS15 K83A* was present at even higher amounts than the endogenous protein in wild-type plants.

**Figure 7.**
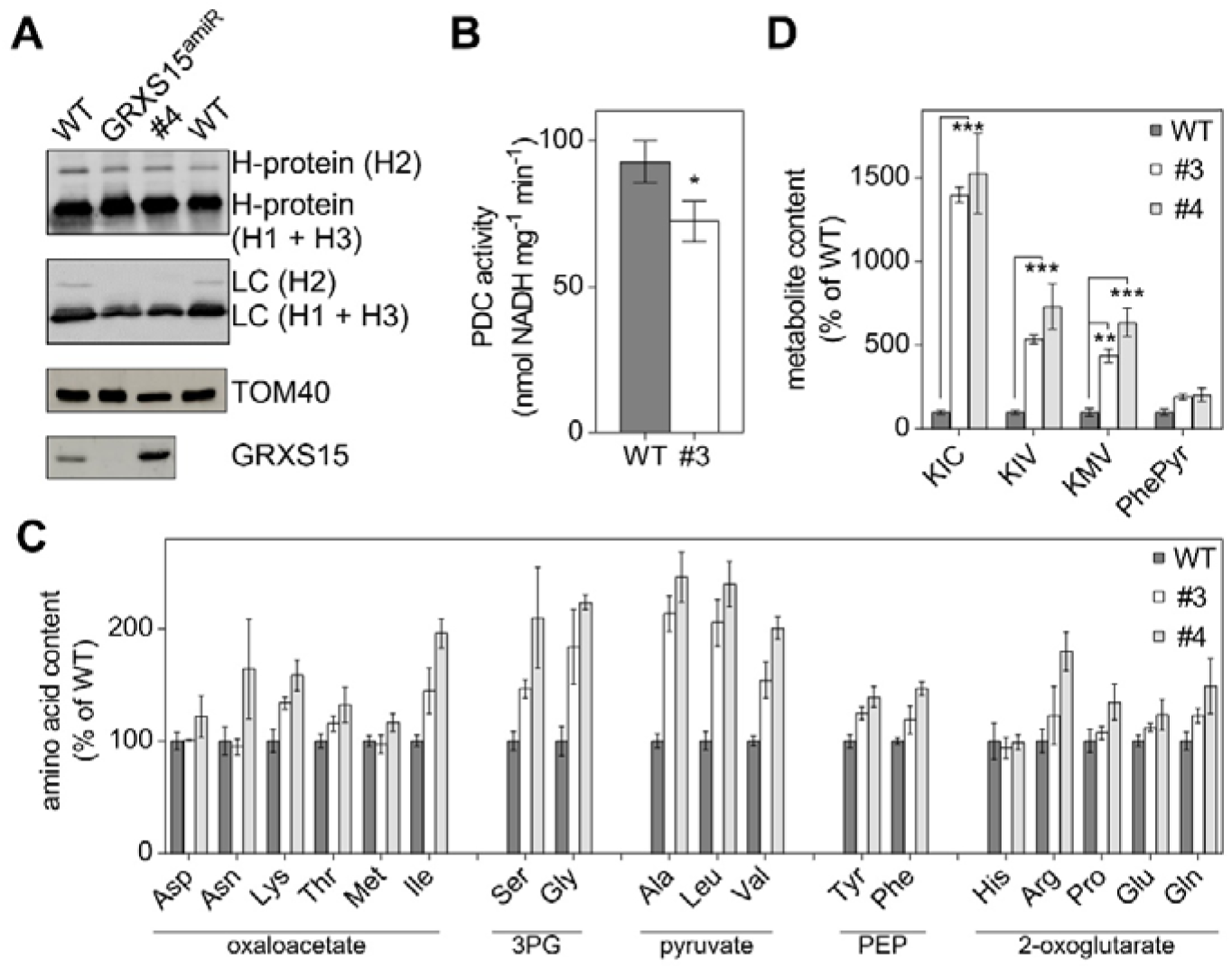
Lipoyl cofactor-dependent enzymes are affected in Arabidopsis *GRXS15 K83A* mutants. **A:** Immunoblot analysis using antibodies against glycine dehydrogenase H-protein (H1-3), lipoyl cofactor (LC) as well as antibodies against TOM40 for a loading control and GRXS15. 15 μg of isolated mitochondria were loaded per lane. **B:** Pyruvate dehydrogenase complex (PDC) activity in isolated mitochondria. Reduction of NAD^+^ was measured in mitochondria isolated from 14-d-old seedlings of WT and the *GRXS15 K83A* line #3 (*n* = 5; means ± SEM). The statistical analysis (one-way ANOVA with post hoc Holm-Sidak comparisons for WT vs. *grxs15* mutant) indicated significant changes; \**P* ≤ 0.05). **C:** Relative abundance of amino acids in 8-d-old seedlings of WT compared *GRXS15 K83A* lines #3 and #4. WT was set to 100% (*n* = 4-5, means ± SEM). Absolute values and statistical analysis are provided in Suppl. Table S1. Amino acids were categorized after their respective common precursor. 3PG = 3-phosphoglycerate, PEP = phosphoenolpyruvate. **D:** Analysis of the breakdown products of leucine, isoleucine and valine – α-ketoisocaproic acid (KIC), α-ketoisovaleric acid (KIV), α-keto-β-methylvaleric acid (KMV) – and phenylpyruvate (PhePyr) in seedlings of WT compared to *GRXS15 K83A* lines #3 and #4. WT was set to 100% (*n* = 4-5; means ± SEM). Absolute values are provided in Suppl. Table S1. The statistical analysis (two-way ANOVA with post hoc Holm-Sidak comparisons for WT vs. *grxs15* mutant) indicated significant changes; \*\**P* ≤ 0.01; \*\*\**P* ≤ 0.001.

To further test whether the accumulation of pyruvate was due to a less active PDC, we measured the activity of the PDC in isolated mitochondria. Interestingly, there was a 22% reduction in activity. While the WT had a PDC activity of 92.7 ± 6.5 nmol NADH mg^-1^ min^-1^ the *GRXS15 K83A* line #3 had a significantly lower activity of only 72.40 ± 6.2 nmol NADH mg^-1^ min^-1^ (Fig. 7B).

The pronounced increase of pyruvate and several TCA intermediates (Fig. 6) may have further effects on downstream metabolites. Given that intermediates of glycolysis and the TCA cycle are hubs for synthesis of amino acids and because mutants defective in PDC subunit E2 show an increase in the pools of nearly all amino acids (Yu et al., 2012), we profiled the abundance of amino acids. Most amino acids were increased in the mutants compared to WT seedlings, with more pronounced increases in line #4 compared to line #3 (Fig. 7C, Supplemental Table S1). Particularly high increases in amino acid abundance of more than 200% were observed for glycine and serine derived from 3-phosphoglycerate, for alanine, leucine and valine all derived from pyruvate, and for isoleucine (Fig. 7C, Supplemental Table S1). The Gly/Ser ratio, indicative of photorespiratory effects, did not show any pronounced change and varied only between 0.33 ± 0.04 for the WT, 0.4 ± 0.1 for line #3 and 0.37 ± 0.12 for line #4.

### Branched-chain amino acid metabolism is strongly impaired in response to diminished GRXS15 activity and lipoyl cofactor availability

Leucine, valine and isoleucine are classified as BCAAs, which share a common degradation pathway that is localized in the mitochondrion. Because the BCAA catabolism pathway involves lipoyl cofactor--dependent BCKDC, the increase in the pools of all three BCAAs may not exclusively result from increased availability of their parent compounds, but also from restricted BCAA degradation capacity. To test this hypothesis, we measured the content of the respective keto acids resulting from deamination of the BCAAs by branched-chain amino acid transaminase (BCAT; Supplemental Fig. 5A). The keto acids α-ketoisocaproic acid (KIC), α-keto-β-methylvaleric acid (KMV) and α-ketoisovaleric acid (KIV) derived from the BCAAs accumulated massively in both *GRXS15 K83A* mutants (Fig. 7D). Here, KIC accumulated in the *GRXS15 K83A* mutants up to 15-fold, resulting in values of 3.5 ± 0.11 pmol (mg FW)^-1^ in line #3 and 3.8 ± 0.6 pmol (mg FW)^-1^ in line #4 compared to 0.25 ± 0.032 pmol (mg FW)^-1^ in the WT. KIV and KMV increased 6 to 7-fold in the *GRXS15 K83A* mutants. These pronounced changes support the hypothesis of decreased BCKDC activity creating a bottleneck in keto acid catabolism (Supplemental Fig. S5A). The higher accumulation of KIC can be accounted for by the preference of BCKDC for the Val derivative (Taylor et al., 2004) resulting in KIV to be metabolized faster and to accumulate less strongly. Despite the presumed bottleneck in catabolism of BCAAs, the *grxs15* mutants did not show enhanced Leu sensitivity (Supplemental Fig. S5B). Similarly, *ivdh* mutants deficient in isovaleryl-CoA dehydrogenase did not display an increased sensitivity to external supply of Leu compared to WT.

## Discussion

### GRXS15 function limits growth

Null mutants of mitochondrial GRXS15 are embryo-defective but can be partially complemented with a mutated GRXS15 protein compromised in its ability to coordinate a [2Fe-2S] cluster (Moseler et al., 2015). The bottleneck in Fe-S coordination results in a dwarf phenotype similar to the phenotype of severe knockdown mutants generated through expression of artificial microRNAs (Supplemental Fig. S1) (Ströher et al., 2016) but how exactly the modification of either activity or abundance of GRXS15 impacts on plant growth and development remained unclear. Less severe knockdown mutants resulting from a T-DNA insertion in the 5’-UTR of *GRXS15* limited the abundance of GRXS15 to about 20% of WT levels, but did not show a macroscopic phenotype beyond early seedling stage under non-stress conditions (Ströher et al., 2016). The growth phenotype of more severe *grxs15* mutants is most apparent in very short roots, which may be linked to the fact that *GRXS15* is strongly expressed in roots, particularly in the maturation and meristematic zone (Belin et al., 2015). The primary function of GRXS15 is assumed to be a role in mitochondrial Fe-S cluster transfer (Moseler et al., 2015; Ströher et al., 2016). This implies that a compromised GRXS15 function potentially may have implications for Fe-S-dependent pathways, including biosynthesis of biotin and Moco, the mETC, and the TCA cycle. While biotin feeding experiments clearly excluded biotin biosynthesis as the limiting factor, the picture was less clear for Moco, which is an essential cofactor for several cytosolic enzymes (Schwarz and Mendel, 2006). Nitrate assimilation, which is dependent on Moco-containing nitrate reductase, initially showed the expected nitrate sensitivity. Measurements of extractable nitrate reductase activity, however, showed no defects. Because, similarly xanthine dehydrogenase and aldehyde oxidases did not show changes in their activities between mutants and the WT, deficiencies in Moco supply can be excluded as a putative metabolic bottleneck in *GRXS15 K83A* mutants. Nitrate sensitivity in *grxs15* mutants leaves us with the conundrum of a different link between mitochondrial functions of GRXS15 and nutrient assimilation, which deserves further investigation in the future.

### GRXS15 does not affect energy balance and ROS levels

Diminished growth correlates with decreased root respiration rates in both, severe *GRXS15^amiR^* knockdown mutants (Ströher et al., 2016) and the weak complementation line #3 investigated in this work (Fig. 4A). Because the mETC contains 12 Fe-S proteins involved in electron transport (Couturier et al., 2015; Meyer et al., 2019) restricted supply of Fe-S clusters would be expected to affect electron flow along the mETC. In humans, it was observed that a patient deficient in mitochondrial glutaredoxin 5 (GLRX5) suffers from decreased abundance and hence activity of complex I (Ye et al., 2010). In yeast, *Δgrx5* mutants displayed a decreased complex II activity, albeit an unaffected protein abundance in this complex (Rodríguez-Manzaneque et al., 2002). In contrast, we found no changes in abundance of any mETC complexes in severe *grxs15* mutants of Arabidopsis (Fig. 4B). Consistently, feeding of mitochondria isolated from *GRXS15 K83A* mutants with succinate revealed that SDH, which contains three different Fe-S clusters (Figueroa et al., 2001), does not constitute a bottleneck in mitochondrial metabolism of *grxs15* mutants. Generally, the respiratory capacity is not affected in the mutants compared to WT, which indicates that supply of Fe-S clusters to components of the mETC is not compromised in *grxs15* mutants. The lower respiratory rate in *GRXS15 K83A* mutants also does not lead to changes in ATP levels. This, however, may also partially be due to decreased consumption of ATP with restricted growth and also the activity of adenylate kinase that contributes to formation of ATP (and AMP) from ADP to buffer the ATP level (De Col et al., 2017). Our overall conclusion to this point is that reduced respiration is likely due to restricted substrate supply rather than assembly of complexes in the mETC and their supply with Fe-S clusters. Restricted supply of reducing equivalents may result from a slowdown of the TCA cycle and also from severely compromised contributions of the electron-transfer flavoprotein/electron-transfer flavoprotein:ubiquinone oxidoreductase (ETF/ETFQO) to ubiquinone reduction (Supplemental Fig. S5). Electrons that enter the mETC via ETF/ETFQO originate from IVDH mediated oxidation of acyl-CoAs as products of BCKDC. The ETF/ETFQO pathway has been shown to contribute significant amounts of electrons in stress situations (Ishizaki et al., 2005; Pires et al., 2016). The concomitant increase in BCKAs and particularly BCAAs may contribute to the dwarf phenotype as disruption in BCAA homeostasis has been shown to lead to pleiotropic effects including growth retardation (Cao et al., 2019).

### GRXS15 affects enzymes and metabolites in the TCA cycle

GRXS15 was detected as part of higher order protein assemblies in a mitochondrial complexome analysis (Senkler et al., 2017). A particularly strong interaction between GRXS15 and mitochondrial isocitrate dehydrogenase 1 (IDH1) was observed in yeast two-hybrid screens with IDH1 as bait and this interaction was subsequently confirmed by bimolecular fluorescence (BiFC) assays (Zhang et al., 2018). Consistent with a suspected role of GRXS15 in IDH1 function, the isocitrate content was decreased significantly in a *grxs15* knockdown mutant, while the relative flux through the TCA cycle increased (Zhang et al., 2018). IDH1 has recently been reported to contain several redox-active thiols that can change their redox state depending on substrate availability for the TCA (Nietzel et al., 2020). The IDH1-GRXS15 interaction thus could point at a possible function of GRXS15 as a thiol-switch operator for regulatory thiols. This is unlikely though, because GRXS15 does not show any reductive activity and only weak oxidative activity (Moseler et al., 2015; Begas et al., 2017). The increase in all analyzed metabolites of the TCA cycle is rather consistent with metabolite patterns found in knockdown mutants of mitochondrial MnSOD, in which increased levels of organic acids correlated with a decrease in ACO activity (Morgan et al., 2008). Aconitase contains a [4Fe-4S] cluster and has frequently been used as an enzymatic marker for defects in Fe-S cluster assembly and transfer in yeast and human cells (Rodríguez-Manzaneque et al., 2002; Bandyopadhyay et al., 2008; Liu et al., 2016). It came as a surprise that ACO was reported to be unaffected in mitochondria of Arabidopsis *grxs15* mutants, both in abundance and activity (Ströher et al., 2016). Consistent with this report, we also found no change in abundance of mitochondrial ACOs, but did find reduced activity (Fig. 5). This decrease in activity may well reflect decreased supply of [4Fe-4S] in line with reports for mutants from non-plant species with defects in mitochondrial Fe-S transfer (Rodríguez-Manzaneque et al., 2002; Liu et al., 2016). In addition, ACOs are prone to oxidative modification by ROS or reactive nitrogen species (Castro et al., 2019) and indeed redox-sensitive thiol residues have been identified on mitochondrial ACOs as putative thiol switches (Nietzel et al., 2020). The absence of any detectable oxidative response, however, provides no lead for further investigation of such speculative redox-dependent regulation of ACO activity under non-stress conditions. The decrease in mitochondrial ACO activity in *GRXS15 K83A* mutants does not explain the most pronounced increase in pyruvate, which accumulates up to five-fold and thus supersedes the accumulation of all other TCA cycle intermediates at least by a factor of two. A knockdown of mitochondrial and cytosolic ACO activities in wild tomato led to a reduction in 2-OG levels but an increase in citrate and isocitrate by 40-50%. A similar change in these organic acids of the TCA cycle were found in a succinate dehydrogenase mutant (Carrari et al., 2003; Huang et al., 2013). The pattern of organic acids in *GRXS15 K83A* mutants is thus clearly different from other TCA cycle mutants. The most pronounced increases in pools of 2-OG and pyruvate compared to WT point at diminished activities of PDC and OGDC instead.

### GRXS15 has a function in protein lipoylation

PDC and OGDC do not contain an Fe-S cluster but rather belong to a class of four dehydrogenase complexes that all involve lipoylated subunits. Lipoylation of mitochondrial proteins is mediated through coordinated action of lipoate-protein ligase, octanoyltransferase, and LIP1 (Ewald et al., 2014). The radical S-adenosylmethionine enzyme LIP1, contains two [4Fe-4S] clusters one of which is required as a substrate, i.e. as sulfur donor to octanoyl-residues (McCarthy and Booker, 2017). Continuous destruction of Fe-S clusters during lipoylation may thus render lipoyl cofactor-dependent-enzymes indirectly sensitive to defects in Fe-S supply. Decreased lipoylation of GDC-H proteins and reduced PDC activity are fully consistent with previous observations on *GRXS15^amiR^* mutants by Ströher et al. (2016). Similar to the Arabidopsis mutants also humans carrying mutations in mitochondrial GLRX5 are deficient in lipoylation of mitochondrial proteins (Baker et al., 2014). A critical restriction through lipoylation deficiency is further supported by increased amounts of pyruvate and 2-OG as well as several other organic acids and amino acids derived from these precursors (Figs. 6 and 7C). Similar increases in pyruvate as well as the accumulation of most amino acids were also shown for Arabidopsis plants with a mutated PDC-E2 subunit resulting in only 30% PDC activity (Yu et al., 2012). A much more pronounced increase of alanine in PDC-E2 mutants than in *GRXS15 K83A* mutants may be attributed to a higher severity of the metabolic bottleneck if PDC activity is down to 30%. Of all metabolites analyzed in this study, the 4 to 15-fold increases of BCKAs in *GRXS15 K83A* mutants were the most pronounced relative changes compared to the WT. The findings that these increases were stronger in more severe mutants, point at BCKDC as a critical bottleneck. The keto acids KIC, KIV and KMV are products of transamination of the BCAAs leucine, isoleucine and valine (Hildebrandt et al., 2015). Further degradation of the keto acids in *GRXS15 K83A* mutants is limited because BCKDC relies on efficient lipoylation of the E2 subunit. Like GDC, PDC and OGDC, BCKDC consists of different subunits, which may not be present in stoichiometric amounts. Recently, Fuchs et al. (2020) reported quantitative data for the abundance of proteins in single mitochondria (Fig. 8). These data indicate low abundance of BCKDC-E2 compared to GDC-H and particularly PDC-E2. Given that all four dehydrogenase complexes rely on the same dihydrolipoyl dehydrogenase subunits, i.e. E3 or L subunits, it is obvious that the relative abundance of subunit proteins will have some impact on assembly of functional complexes. With the assumption that all different E2 and H subunits compete with each other and with the same chance of getting lipoylated, the absolute number of lipoylated PDC-E2 proteins is expected to be higher than that of BCKDC-E2 proteins. If even non-lipoylated E2 or H formed complexes with E3 or L, very little functional BCKDC could be formed. Furthermore, the very low copy number of BCKDC-E1 subunits compared to BCKDC-E2 implies that under lipoyl cofactor-limiting conditions, E1 subunits are more likely to form complexes with non-lipoylated and hence non-catalytic E2. *Vice versa*, the few E2 copies that do get lipoylated may not be those that assemble with E1 subunits to form active complexes. Selective transcriptional upregulation of several nominally lipoylated subunits in *GRXS15^amiR^* as reported by Ströher et al. (2016) would additionally increase the imbalance and tighten the metabolic bottleneck even further.

**Figure 8.**
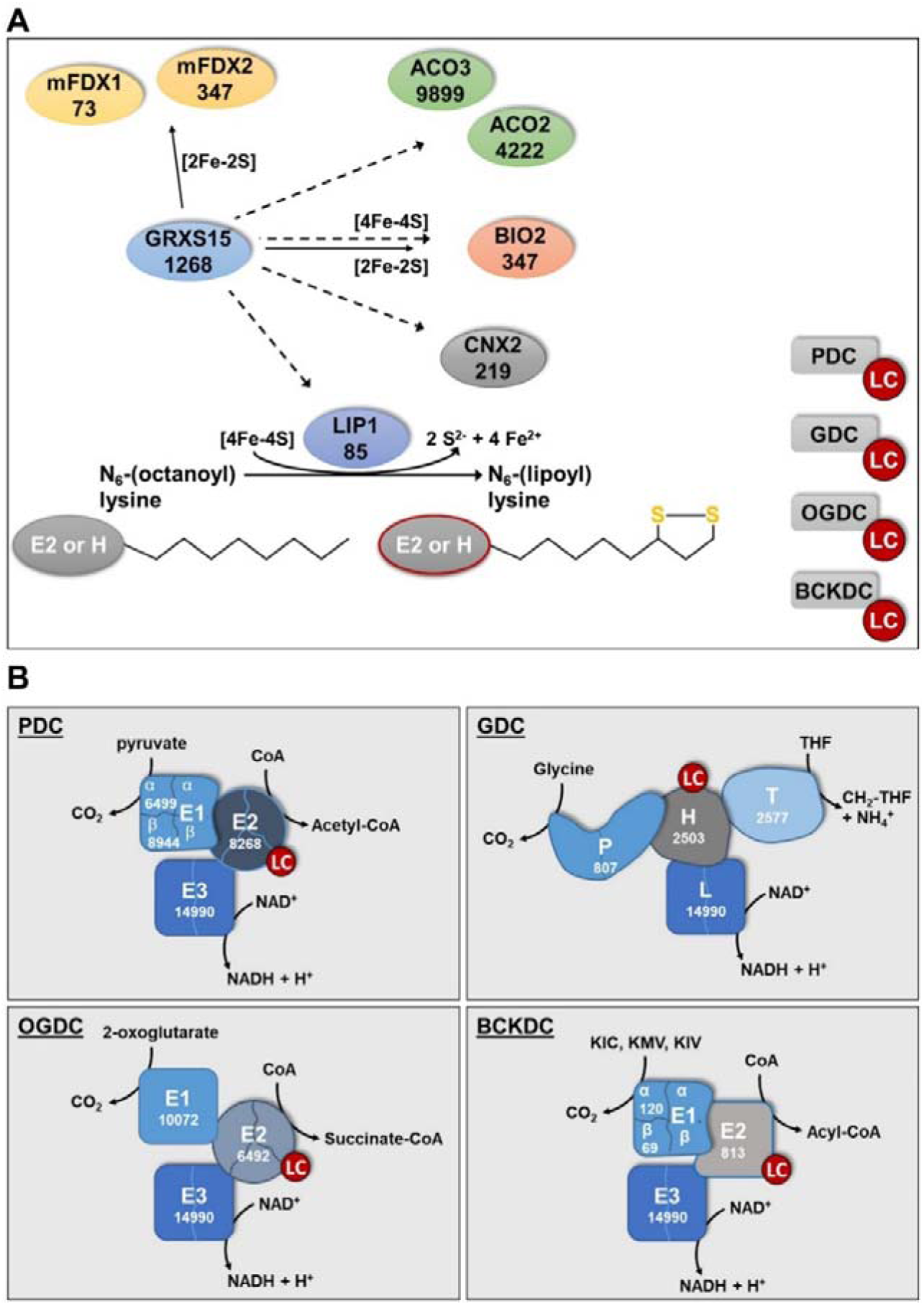
Lipoylation of mitochondrial proteins depends on GRXS15. **A:** Distribution of Fe-S clusters in Arabidopsis mitochondria to soluble Fe-S proteins and lipoylation of proteins via lipoyl synthase (LIP1). Putative transfer of Fe-S clusters is indicated by solid arrows for [2Fe-2S] and dashed arrows for [4Fe-4S]. Intermediate complexes of Fe-S transfer and assembly of [4Fe-4S] clusters are not shown. mFDX1/2: mitochondrial ferredoxin 1/2; ACO2/3; aconitase 2/3; BIO2: biotin synthase 2; CNX2: GTP-3’,8-cyclase PDC: pyruvate decarboxylase complex; OGDC: 2-oxoglutarate dehydrogenase complex; GDC: glycine decarboxylase complex; BCKDC: branched-chain α-keto acid dehydrogenase complex; LC: Lipoyl cofactor. Numbers give the estimated copy number of the respective proteins according to Fuchs et al. (2020). **B:** Abundance of subunits in the four mitochondrial dehydrogenase complexes PDC, GDC, OGDC and BCKDC according to Fuchs et al. (2020). The copy number for the H subunit of GDC is only for the isoform H2, because the nominally more abundant isoforms H1 and H3 (see Fig. 7A) were not identified by Fuchs et al. (2020). E3 and L subunits are formed by the closely related and highly similar proteins mtLPD1 (4876 copies) and mtLPD2 (10114 copies), The total of both isoforms is given but it should be noted that a preference of GDC for mtLPD1 and of the other three complexes for mtLPD2 has been proposed (Lutziger and Oliver, 2001). Deficiencies of these enzymes generates metabolic bottlenecks and causes an increase of their respective substrates and particularly for PDC and OGDC also a severe limit in carbon supply to the TCA cycle.

LIP1 was estimated to be present with 85 copies in a single mitochondrion compared to 4200 copies of ACO2 and 9900 copies of ACO3 (Fig. 8) (Fuchs et al., 2020). In the absence of any other evidence all apoproteins have a similar likelihood of receiving a [4Fe-4S] cluster. The few LIP1 proteins will have a low chance of receiving a cluster if efficient supply of [2Fe-2S] clusters by GRXS15 further upstream in the Fe-S cluster transfer machinery is compromised. With the need for two [4Fe-4S] clusters of which one has to be replaced after each catalytic cycle the bottleneck is bound to become even more severe than in enzymes that use their Fe-S clusters only for electron transfer reactions.

## Conclusion

We show that compromising the ability of GRXS15 to coordinate and transfer [2Fe-2S] clusters results in severe defects only in enzymes relying on the prosthetic group lipoamide. These results are in agreement with findings by Ströher et al. (2016) who reported diminished lipoylation of proteins in *GRXS15^amiR^* lines and hypothesized that diminished respiration and the short root mutant phenotype could be a consequence of the incomplete lipoyl cofactor loading of important TCA cycle enzymes. Here we expand and specify the picture, by systematically probing for metabolic bottlenecks in mitochondrial pathways that rely on the supply with Fe-S clusters. While changes in a several metabolites were found, the primary defects can be assigned to merely the four mitochondrial dehydrogenase complexes all of which contain a lipoylated subunit. Those results emphasize the importance of LIP1 as a major sink for Fe-S clusters, which becomes manifest if GRXS15-mediated Fe-S cluster transfer between the assembly machinery and receiving apoproteins is restricted. The fact that most other Fe-S-dependent pathways are not seriously affected by deficiencies in *GRXS15 K83A* complementation lines may be explained by the effective relative abundance of different proteins in mitochondria. We propose that an increased demand for Fe-S as sulfur donor combined with the very low abundance of LIP1 leads to the manifestation of a potentially lethal bottleneck. The phenotype highlights the importance of an accurate maintenance of protein amounts and appropriate stoichiometries for normal mitochondrial function.

## Material and Methods

### Plant Material & Methods

Previously described *Arabidopsis thaliana* complementation lines *grxs15-3 UBQ10_pro_:GRXS15 K83A* (Moseler et al., 2015) and the knock-down line *GRXS15^amiR^* (Ströher et al., 2016) as well as *atm3-1* and *atm3-4* (Teschner et al., 2010) were used in this study. *A. thaliana* ecotype Col-0 (([L.] Heyn.) segregated from the T-DNA line *grxs15-3*) was used as WT. Unless stated differently, surface-sterilized seeds were grown on vertical plates containing nutrient medium (Meyer and Fricker, 2000) with 0.1% (w/v) sucrose solidified with 0.8% (w/v) agar under long-day conditions with a diurnal cycle of 16 h light at 22°C and 8 h dark at 18°C. The light intensity was 75 μE m^−2^ s^−1^ and 50% air humidity.

Germination rate was scored by observing radical emergence in seeds plated on vertical culture plates using a stereomicroscope (Leica M165 FC). Root growth was documented photographically on vertical culture plates containing 0.8% (w/v) phytagel and 0.1% (w/v) sucrose. Five and 8 d after stratification, root length was documented and measured using Adobe Illustrator CS5.1.

Influence of the nitrogen source on root length was analyzed on plates containing 5 mM KNO_3_ or 2.5 mM (NH_4_)_2_SO_4_, 2.5 mM KH_2_PO_4_, 2 mM MgSO_4_, 2 mM CaCl_2_, 50 μM Fe-EDTA, 70 µM H_3_BO_4_, 14 µM MnCl_2_, 0.5 µM CuSO_4_, 1 µM ZnSO_4_, 0.2 µM NaMoO_4_, 10 µM NaCl, 0.01 µM CoCl_2_, 0.8% (w/v) phytagel and 0.1% (w/v) sucrose, pH 5.8. To check for possible effects of counter anions, (NH_4_)_2_SO_4_ was replaced by NH_4_Cl and grown otherwise exactly under the same conditions.

### Blue Native Page

Mitochondrial samples were solubilized in 1% (w/v) n-dodecyl β-D-maltoside and subjected to Blue-Native-PAGE as described previously (Meyer et al., 2011; Kühn et al., 2015).

### Isolation of mitochondria

Arabidopsis mitochondria were purified from 2- or 4-week-old seedlings as described before (Sweetlove et al., 2007) with slight modifications. All steps were performed on ice or at 4°C. Seedlings were homogenized using mortar and pestle and the homogenate was filtered (Miracloth; Merck Millipore) before cellular debris was pelleted by centrifugation for 5 min at 1,200 *g*. The supernatant was centrifuged for 20 min at 18,000 *g*, and the pellet of crude mitochondria was gently resuspended in wash buffer (0.3 M sucrose, 0.1% (w/v) BSA and 10 mM TES, pH 7.5) and centrifuged for 5 min at 1,200 *g*. The supernatant was transferred into a new tube and centrifuged for 20 min at 18,000 *g*. The pellet was gently resuspended in final wash buffer (0.3 M sucrose, 10 mM TES, pH 7.5), loaded directly on a 0–6% Percoll gradient and centrifuged for 40 min at 40,000 *g*. Mitochondria were transferred into a new tube and washed three times with final wash buffer (0.3 M sucrose, 10 mM TES pH 7.5).

### Respiration Assays

Oxygen consumption of intact Arabidopsis roots and isolated mitochondria was measured in Oxytherm Clark-type electrodes (Hansatech; www.hansatech-instruments.com) as described before (Wagner et al., 2015). Whole roots from seedlings vertically grown on agar plates were cut below the hypocotyl-root junction and assayed in a volume of 1.2 mL containing 5 mM KCl, 10 mM MES, and 10 mM CaCl_2_, pH 5.8, and after addition of 4 mM KCN and 0.2 mM pGal.

O_2_ consumption of isolated mitochondria was measured in a volume of 1 mL containing M mannitol, 10 mM TES-KOH pH 7.5, 5 mM KH_2_PO_4_, 10 mM NaCl, 2 mM MgSO_4_ and 0.1% (w/v) bovine serum albumin. O_2_ consumption rate was measured before (blank) addition of mitochondria and after addition of mitochondria or respective substrate (state II; succinate (10 mM succinate, 0.25 mM ATP) or pyruvate/malate (10 mM pyruvate, 10 mM malate, 0.3 mM NAD and 0.1 mM thiamine pyrophosphate), state III; ADP (50 μM ADP). Additionally, O_2_ consumption rate was analyzed after ADP consumption (state IV) and after addition of 10 μM carbonyl cyanide m-chlorophenylhydrazone (CCCP).

### Histological detection of reactive oxygen species

For detection of increased H_2_O_2_ production, leaves were stained with DAB (3, 3-diaminobenzidine) (Thordal-Christensen et al., 1997). Leaves were vacuum-infiltrated in a solution containing 0.1 mg mL^-1^ DAB, 50 mM potassium phosphate buffer pH 7.6 and 0.1% (v/v) Silwet L-77. After infiltration, the leaves were incubated for 20-24 h in the dark and destained by lactic acid:glycerol:EtOH (1:1:3) for 30 min at 70°C.

For histochemical staining of superoxide, NBT (nitro blue tetrazolium) was used (Hoffmann et al., 2013). Leaves were vacuum-infiltrated in a solution containing 0.1 mg mL^-1^ NBT, 50 mM potassium phosphate buffer pH 7.6 and 0.1% (v/v) Silwet L-77. After infiltration the leaves were incubated for 30 min in the dark and destained by lactic acid:glycerol:EtOH (1:1:3) for 30 min at 70°C.

### Determination of metabolite levels via HPLC

Aliquots (45-55 mg) of freshly ground plant tissue were used for absolute quantification of amino acid, α-keto acid and organic acid content each.

Free amino acids and α-keto acids were extracted with 0.5 mL ice-cold 0.1 M HCl in an ultrasonic ice-bath for 10 min. Cell debris and insoluble material were removed by centrifugation for 10 min at 25,000 *g*. For the determination of α-keto acids, 150 µL of the resulting supernatant were mixed with an equal volume of 25 mM OPD (o-phenylendiamine) solution and derivatised by incubation at 50°C for 30 min. After centrifugation for 10 min, the derivatised keto acids were separated by reversed phase chromatography on an Acquity HSS T3 column (100 mm x 2.1 mm, 1.7 µm, Waters) connected to an Acquity H-class UPLC system. Prior separation, the column was heated to 40°C and equilibrated with 5 column volumes of solvent A (0.1% (v/v) formic acid in 10% (v/v) acetonitrile) at a flow rate of 0.55 mL min^-1^. Separation of keto acid derivatives was achieved by increasing the concentration of solvent B (acetonitrile) in solvent A (2 min 2% B, 5 min 18% B, 5.2 min 22% B, 9 min 40% B, 9.1 min 80% B and hold for 2 min, and return to 2% B in 2 min). The separated derivatives were detected by fluorescence (Acquity FLR detector, Waters, excitation: 350 nm, emission: 410 nm) and quantified using ultrapure standards (Sigma). Data acquisition and processing were performed with the Empower3 software suite (Waters). Derivatisation and separation of amino acids was performed as described by (Yang et al., 2015).

Total organic acids were extracted with 0.5 mL ultra-pure water for 20 min at 95°C. Organic acids were separated using an IonPac AS11-HC (2 mm, Thermo Scientific) column connected to an ICS-5000 system (Thermo Scientific) and quantified by conductivity detection after cation suppression (ASRS-300 2 mm, suppressor current 95-120 mA). Prior separation, the column was heated to 30°C and equilibrated with 5 column volumes of solvent A (ultra-pure water) at a flow rate of 0.38 mL min^-1^. Separation of anions and organic acids was achieved by increasing the concentration of solvent B (100 mM NaOH) in buffer A (8 min 4% B, 18 min 18% B, 25 min 19% B, 43 min 30% B, 53 min 62% B, 53.1 min 80% B for 6 min, and return to 4% B in 11 min). Soluble sugars were separated on a CarboPac PA1 column (Thermo Scientific) connected to the ICS-5000 system and quantified by pulsed amperometric detection (HPAEC-PAD). Column temperature was kept constant at 25°C and equilibrated with five column volumes of solvent A (ultra-pure water) at a flow rate of 1 mL min^-1^. Baseline separation of carbohydrates was achieved by increasing the concentration of solvent B (300 mM NaOH) in solvent A (from 0 to 25 min 7.4% B, followed by a gradient to 100% B within 12 min, hold for 8 min at 100% B, return to 7.4% B and equilibration of the column for 12 min). Data acquisition and quantification was performed with Chromeleon 7 (Thermo Scientific).

### Aldehyde oxidase and xanthine dehydrogenase assay

Aldehyde oxidase (AO) and xanthine dehydrogenase (XDH) assays were performed similar as described previously by Koshiba et al. (1996) and Hesberg et al. (2004). For determination of AO and XDH activities Arabidopsis seedlings were homogenized in extraction buffer (0.1 M potassium phosphate buffer pH 7.5, 2.5 mM EDTA and 5 mM DTT) and centrifuged for 10 min at 16,000 *g* and 4°C. Enzyme activities of AO and XDH in the resulting supernatant were detected after native PAGE by activity staining. Activity of AO was developed in a reaction mixture containing 0.1 M potassium phosphate buffer pH 7.5, 1 mM 1-naphthaldehyde, 1 mM indole-3-carboxaldehyde, 0.1 mM phenazine methosulfate (PMS), and mM MTT (3-(4,5-dimethylthiazol-2-yl)-2,5-diphenyltetrazolium bromide) at RT. Activity of XDH was analyzed with a staining solution of 1 mM hypoxanthine, 1 mM MTT and 0.1 mM PMS in 250 mM Tris-HCl, pH 8.5.

### Nitrate Reductase assay

Nitrate reductase (NR) assay was performed as described previously (Scheible et al., 1997) with slight modifications. Leaves were homogenized in extraction buffer (50 mM MOPS, pH 7.0, 50 mM KCl, 5 mM Mg-acetate, 1 mM CaCl_2_, 2 mM Na-citrate and 1 mM DTT) and centrifuged for 10 min at 20,000 *g* and 4°C. NR activity was measured in a reaction mixture containing 50 mM MOPS, pH 7.0, 50 mM KCl, 5 mM Mg-acetate, 1 mM CaCl_2_, 10 mM KNO_3_ and 0.4 mM NADH. At consecutive time points, 150 µL aliquots were removed from the mixture and the reaction was stopped by adding 54 mM zinc acetate and 37.5 µM PMS. Thereafter, 0.475% (v/v) sulfanilamide in 1 N HCl and 0.005% (v/v) N-(1-naphthyl)-ethylenediamine was added. Samples were allowed to stand for 15 min at RT in the dark and the absorbance of the produced azo-dye was measured at 540 nm.

### Aconitase assay

Aconitase activity was analyzed in a coupled assay measuring NADPH formation by monitoring the increase in absorbance at 340 nm using a plate reader (CLARIOstar^®^; BMG). The reaction mixture contained 50 mM HEPES pH 7.8, 2.5 mM NADP^+^, 5 mM MnCl_2_, 0.1% (v/v) Triton X-100 and 0.05 U isocitrate dehydrogenase. The mixture was allowed to come to equilibrium after addition of protein extract. The reaction was started by adding 8 mM cis-aconitic acid.

### Pyruvate dehydrogenase complex assay

To estimate the activity of pyruvate dehydrogenase complex, mitochondria were isolated as described previously and reduction of NAD^+^ was measured at 340 nm in a reaction mixture containing ∼10 µg mitochondria in 100 mM MOPS pH 7.4, 1 mM CaCl_2_, 1 mM MgCl_2_, 4 mM cysteine, 0.45 mM thiamine pyrophosphate, 0.18 mM Coenzyme A, 3 mM NAD^+^ and 0.1% (v/v) Triton X-100. The reaction was started with 7.5 mM pyruvate.

### Fatty Acid Methyl Ester (FAME) Measurement

The analysis of fatty acids was performed by quantification of their respective fatty acid methyl esters (FAMEs) via gas chromatography coupled with a flame ionization detector as described before (Browse et al., 1986). 1 mL 1 N HCl in MeOH was added to 5 seeds or ∼50 mg homogenized seedlings as well as 5 µg pentadecanoic acid as internal standard. Samples were incubated at 80°C for 2 h (seeds) or 30 min (seedlings). After cooling down, 1 mL 0.9% (w/v) NaCl and 1 mL hexane were added. Samples were mixed vigorously and centrifuged with 1,000 *g* for 3 min. The hexane phase was transferred to a GC vial. FAMEs were quantified using pentadecanoic acid as internal standard.

### Western Blotting

For protein blot analysis, total cell extract or purified organelles were heated for 5 min and separated on standard SDS/PAGE gels. Proteins were transferred to a membrane (BioTrace PVDF Transfer Membrane; Pall Corporation) and labeled with antibodies (Streptavidin HRP: ab7403 Abcam; lipoic acid: ab58724, aconitase: see Bernard et al. (2009). The GRXS15 antibody was a kind gift of Nicolas Rouhier (Nancy) and the H protein antibody a kind gift of Olivier Keech (Umea). The TOM40 antibody was a kind gift of Jim Whelan (Melbourne). Immunolabeling was detected by chemiluminescence by using secondary horseradish peroxidase-conjugated antibodies and Pierce ECL Western Blotting Substrate.

### Fluorescence microscopy

Fluorescent plants were selected using a stereomicroscope (Leica M165 FC) equipped with a GFP filter.

A confocal laser scanning microscope (Zeiss LSM 780, attached to an Axio Observer.Z1; Carl Zeiss Microscopy) and a ×40 (C-Apochromat, 1.20 numerical aperture, water immersion) or ×63 lens (Plan-Apochromat, 1.40 numerical aperture, oil immersion) was used for confocal imaging. For ratiometric analyses of mitochondrial localized roGFP2-hGrx1 (Albrecht et al., 2014) or roGFP2-Orp1 (Nietzel et al., 2019), lines with similar expression levels in WT and mutants were selected. For both sensors, roGFP2 was excited at 405 and 488 nm. For both excitation wavelengths, roGFP2 fluorescence was collected with a bandpass filter of 505-530 nm.

The cytosolic ATeam 1.03-nD/nA was excited at 458 nm and emission of CFP (mseCFP) and Venus (cp173-mVenus) was collected at 499-544 nm and 579-615 nm, respectively. Background signal was subtracted before ratiometric analysis.

For all emissions, intensities from four scans were averaged. Ratiometric analysis was performed using a custom-written MATLAB script (Fricker, 2016) using x,y noise filtering and fluorescence background subtraction.

### Statistical analysis

Statistics and error bars were applied for independent experiments with at least three biological replicates using the program GraphPad Prism 6.

## Supplemental Data

**Supplemental Figure S1:**
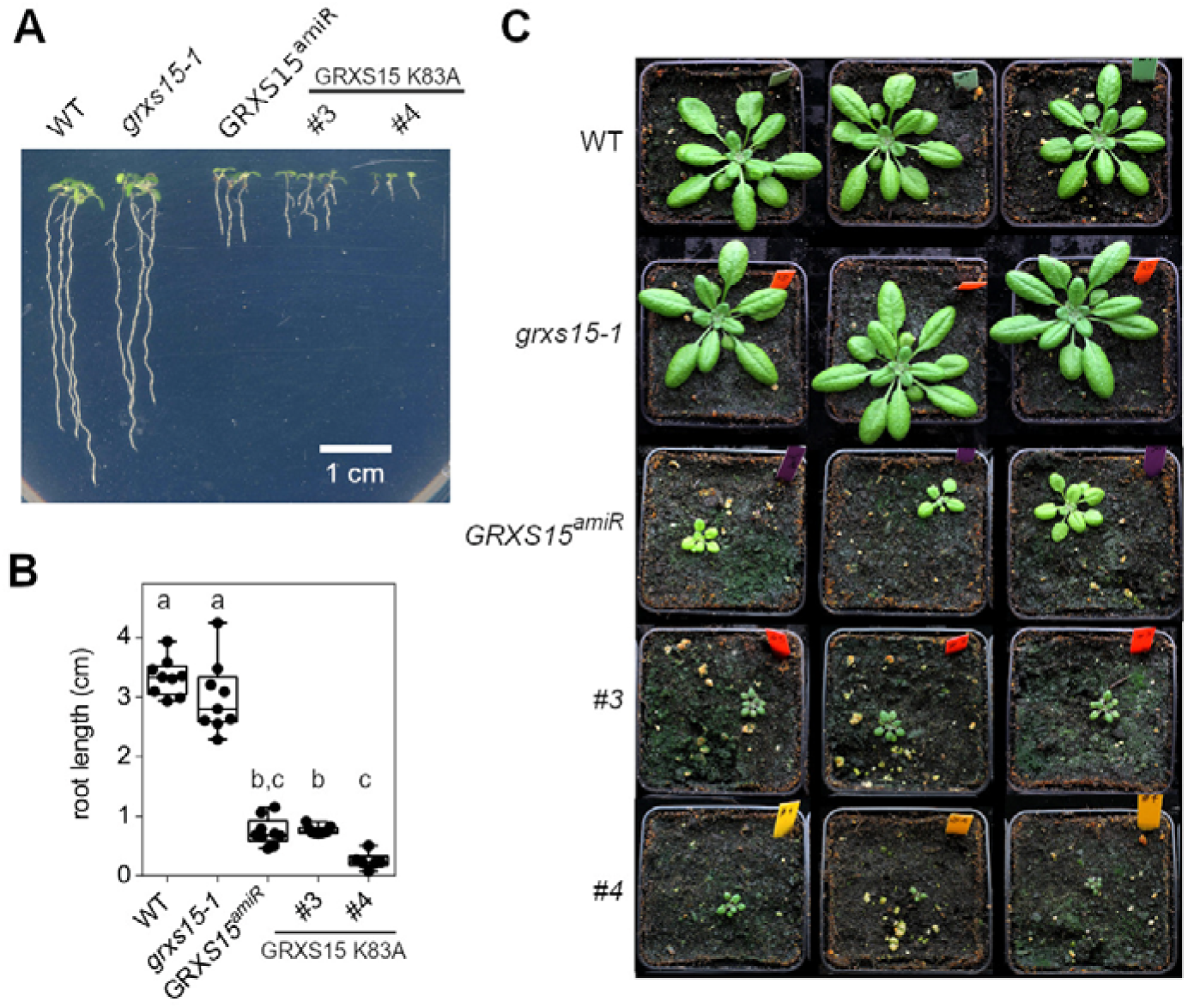
Arabidopsis mutants affected in GRXS15 function develop a dwarf phenotype. **A, B:** Growth of different *grxs15* mutants (*grxs15-1*, *GRXS15^amiR^*, GRXS15 K83A lines #3 and #4) and wild-type (WT) seedlings on vertical plates with 0.8% agar under long-day conditions. Seedlings were documented and quantitatively analyzed for their root length 10 days after germination. (*n* = 6-9; box plot shows means with whiskers indicating min and max values). Different letters indicate significant differences between the different lines; P ≤ 0.05; (one-way ANOVA). **C:** Phenotypes of soil-grown plants after five weeks under long-day conditions (16 h light, 19°C, 8 h dark, 17°C; 50% rh).

**Supplemental Figure S2.**
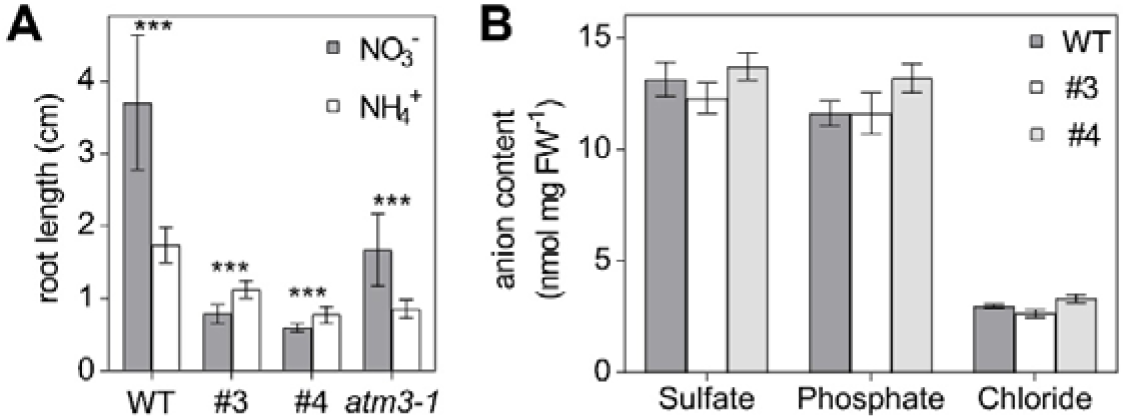
Moco enzymes and anions are not affected in Arabidopsis *GRXS15 K83A* mutants. **A:** Primary root length of *GRXS15 K83A* lines #3 and #4 as well as *atm3-1* mutant seedlings compared to WT grown on vertical plates containing 5 mM KNO_3_ or 5 mM NH_4_Cl as N-source. Seedlings were grown for 9 d under long-day conditions (*n* = 35; means ± SD). Student’s t-Test analysis showed significant differences between nitrate and ammonium treatment for each genotype (***: *P* ≤ 0.001). **B:** Amount of sulfate, phosphate and chloride in Arabidopsis WT and line #3 and #4 seedlings (*n* = 4; means ± SEM). The statistical analysis (two-way ANOVA with post hoc Holm-Sidak comparisons for WT vs. *grxs15*) indicated no significant (*P* ≤ 0.05) change.

**Supplemental Figure S3.**
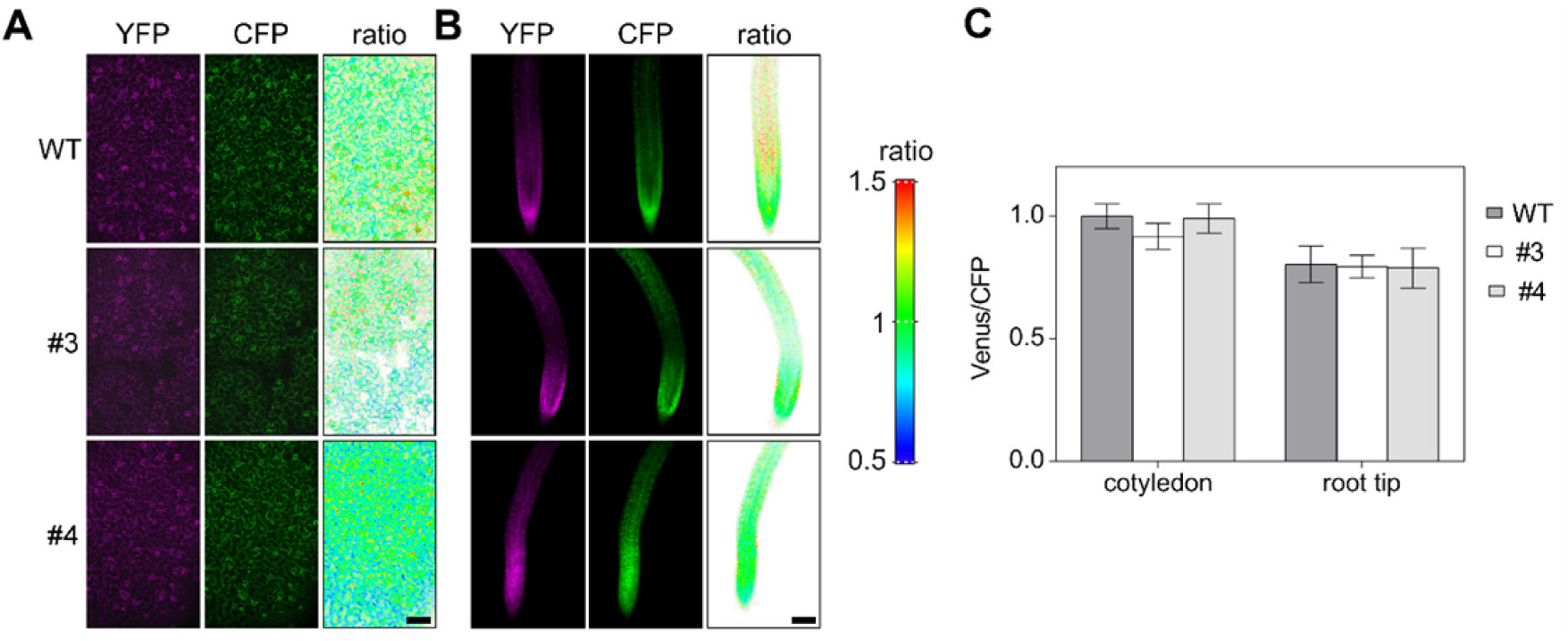
I*n vivo* monitoring of ATP levels in the cytosol of Arabidopsis *GRXS15 K83A* mutants. ATeam1.03-nD/nA was stably expressed under a *35S* promoter in the cytosol of WT and *GRXS15 K83A* lines #3 and #4 and analyzed in cotyledons (A) and roots (B) for fluorescence intensities of Venus and CFP. Bars, 100 µm. (C) Venus/CFP fluorescence ratios calculated from fluorescence images of cotyledons and root tips of 7-d-old seedlings from two independent reporter lines for each genetic background (*n* = 10; means ± SD).

**Supplemental Figure S4.**
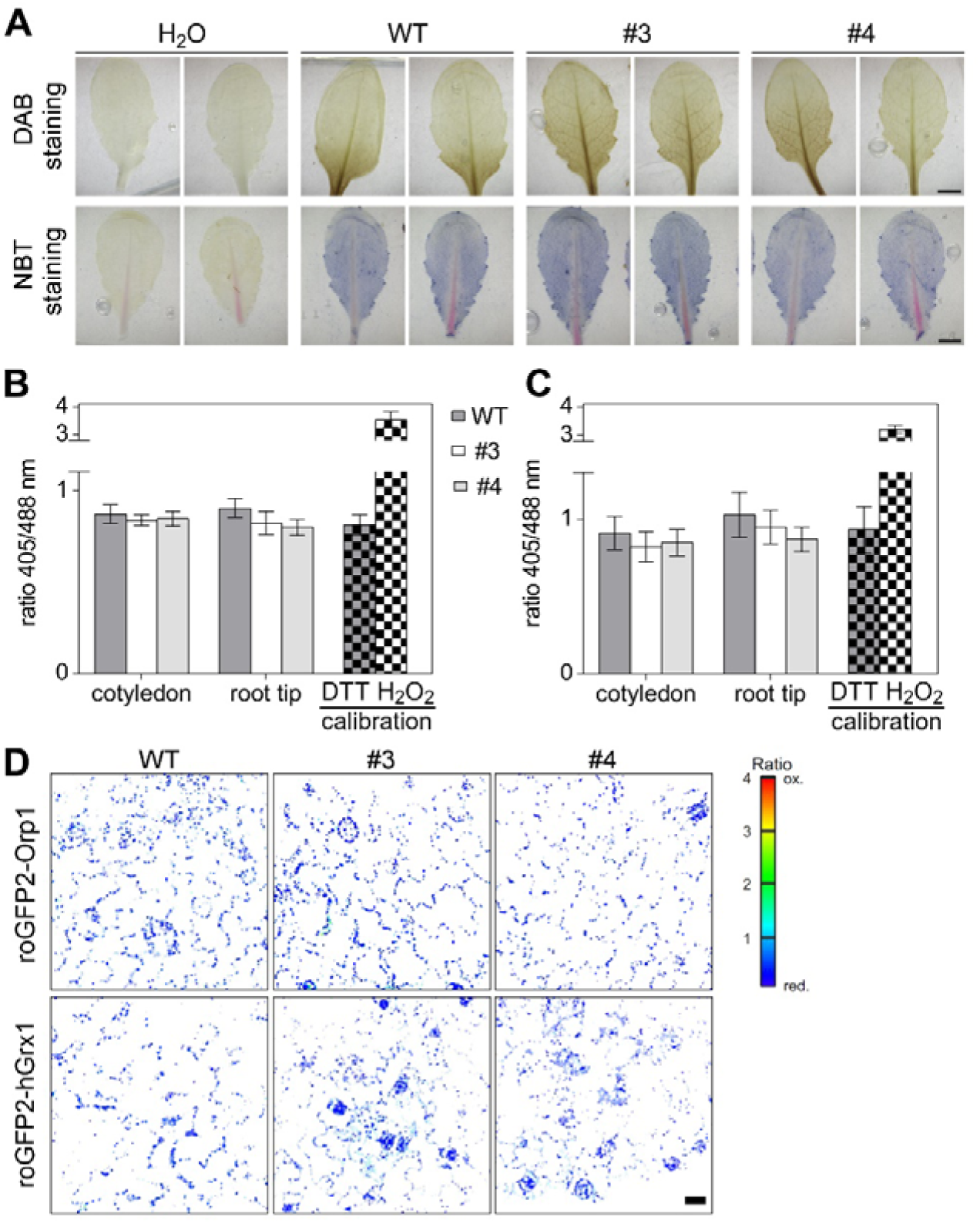
Analysis of the oxidation state of the Arabidopsis *grxs15* mutants. **A:** Representative images showing DAB (upper) and NBT (lower) staining for detection of increased ROS production in leaves. Wild-type plants and mutants were grown for four weeks under long-day growth conditions. Bars, 0.5 cm. *n* = 7-8. **B:** Ratiometric analysis of the H_2_O_2_-sensitive fluorescent reporter roGFP2-Orp1. 7-d-old seedlings of WT and *GRXS15 K83A* lines #3 and #4 expressing mitochondrial roGFP2-Orp1 were analyzed for the redox state of the sensor in cotyledons and root tips. For estimation of the dynamic range of the sensor, wild-type seedlings were incubated in 10 mM DTT (grey squared) or 10 mM H_2_O_2_ (white squared) and fluorescence of roGFP2 in the hypocotyl was analyzed. Ratios were calculated from fluorescence images of cotyledons and root tips of 7-d-old seedlings from two independent reporter lines for each genetic background (*n* = 10; means ± SD). **C:** Ratiometric analysis of the *E*_GSH_–sensitive fluorescent reporter roGFP2-hGrx1 in mitochondria. Ratiometric analysis was performed with 7-d-old seedlings of WT and *GRXS15 K83A* lines #3 and #4 expressing mitochondrial roGFP2-hGrx1 by CLSM. For estimation of the dynamic range of the sensor, wild-type seedlings were incubated in 10 mM DTT (grey squared) or 10 mM H_2_O_2_ (white squared) and fluorescence of roGFP2 in the root tips was analyzed. Ratios were calculated from fluorescence images of cotyledons and root tips of 7-d-old seedlings from two independent reporter lines for each genetic background (*n* = 10; means ± SD). **D:** Representative false color images of cotyledons of 7-d-old seedlings show the oxidation state of roGFP2-Orp1 or roGFP2-hGrx1 targeted to the mitochondrial matrix in WT and *GRXS15 K83A* lines #3 and #4. Bar, 20 µm.

**Supplemental Figure S5.**
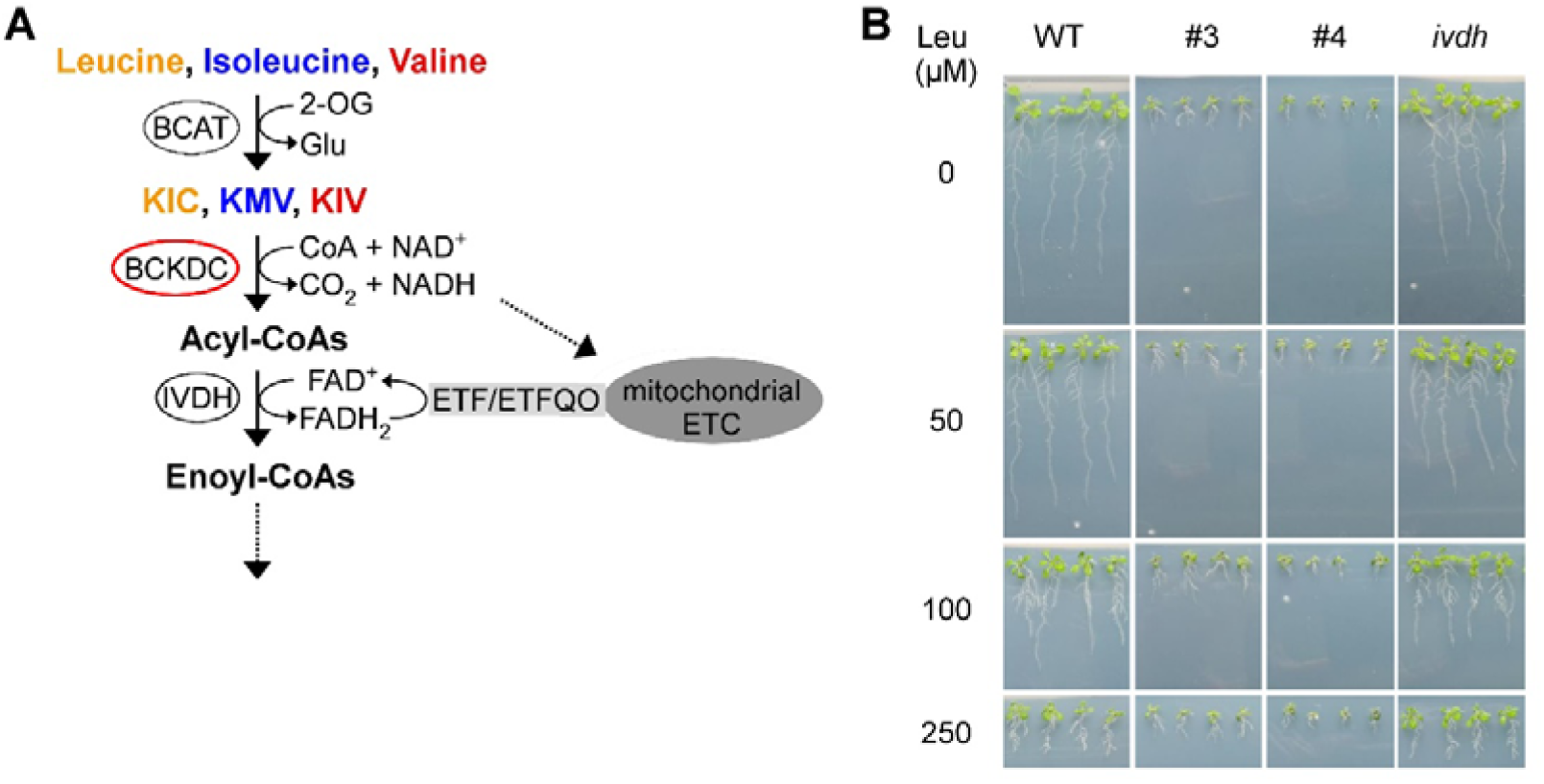
Catabolism of branched-chain amino acids in Arabidopsis seedlings. **A:** The branched-chain amino acids leucine, isoleucine and valine are deaminated by branched-chain aminotransferase (BCAT), which uses largely 2-oxoglutarate (2-OG) forming the branched-chain keto acids α-ketoisocaproic acid (KIC), α-keto-β-methylvaleric acid (KMV) and α-ketoisovaleric acid (KIV) as well as glutamate. The keto acids are further degraded by branched-chain keto acid dehydrogenase (BCKDC), which catalyzes the oxidative decarboxylation producing thereby acyl-CoA and NADH. Isovaleryl-CoA dehydrogenase (IVDH) catalyzes the third step providing electrons to the electron transport chain (ETC) via electron transfer flavoprotein (ETF)/electron transfer flavoprotein ubiquinone oxidoreductase (ETFQO) (modified after Peng et al. (2015)). **B:** Leucine sensitivity of WT, *GRXS15 K83A* lines #3 and #4 and *ivdh* mutants. 4-d-old seedlings were transferred to plates containing the respective leucine amount and were analyzed after 7 d.

**Supplementary Table S1.** Content of amino acids and keto acids of Arabidopsis WT and G*RXS15 K83A* lines #3 and #4. The statistical analysis (two-way ANOVA with post hoc Holm-Sidak comparisons for WT vs. *grxs15* mutant) indicated significance levels; \**P* ≤ 0.1, \*\**P* ≤ 0.01; \*\*\**P* ≤ 0.001; ns: not significant.

| Amino acid | amount of amino acid (pmol (mg FW)<sup>-1</sup>); mean ± SEM (% compared to WT; significance level) |  |  |
| --- | --- | --- | --- |
|  | WT | #3 | #4 |
| Ala | 96.66 ± 6.2 | 206.49 ± 15.25 (213.6; ***) | 237.99 ± 21.35 (246.2; ***) |
| Leu | 6.48 ± 0.53 | 13.32 ± 1.36 (205.5; ***) | 15.55 ± 1.3 (239.8; ***) |
| Gly | 36.79 ± 4.84 | 67.70 ± 12.19 (184.0; ***) | 82.26 ± 2.38 (223.6; ***) |
| Ser | 111.74 ± 9.6 | 163.90 ± 9.11 (146.7; *) | 234.56 ± 50.21 (209.9; ***) |
| Val | 16.48 ± 0.72 | 25.45 ± 2.63 (154.5; *) | 33.03 ± 1.64 (200.5; ***) |
| Ile | 5.45 ± 0.29 | 7.89 ± 1.1 (144.7; *) | 10.70 ± 0.7 (196.3; ***) |
| Arg | 9.96 ± 1.07 | 12.22 ± 2.56 (122.7; ns) | 17.94 ± 1.72 (180.1; ***) |
| Asn | 76.35 ± 9.27 | 72.38 ± 5.36 (94.8; ns) | 125.24 ± 33.73 (164.0; **) |
| Lys | 6.61 ± 0.65 | 8.85 ± 0.36 (133.9; ns) | 10.49 ± 0.9 (158.6; *) |
| Gln | 213.64 ± 17.12 | 262.03 ± 13.91 (122.7; ns) | 317.91 ± 52.1 (148.8; *) |
| Phe | 6.32 ± 0.18 | 7.49 ± 0.8 (118.6; ns) | 9.26 ± 0.36 (146.6; ns) |
| Tyr | 1.50 ± 0.09 | 1.87 ± 0.08 (124.9; ns) | 2.09 ± 0.14 (139.2; ns) |
| Pro | 31.51 ± 3.24 | 33.73 ± 1.87 (107.0; ns) | 42.39 ± 5.11 (134.5; ns) |
| Thr | 58.94 ± 3.66 | 68.05 ± 4.03 (115.5; ns) | 77.90 ± 9.11 (132.2; ns) |
| Glu | 652.08 ± 32.85 | 730.33 ± 22.23 (112.0; ns) | 805.71 ± 90.0 (123.6; ns) |
| Asp | 199.61 ± 15.02 | 200.89 ± 1.35 (100.6; ns) | 243.28 ± 37.33 (121.9; ns) |
| Met | 1.48 ± 0.07 | 1.44 ± 0.12 (97.3; ns) | 1.72 ± 0.12 (116.7; ns) |
| His | 13.46 ± 2.15 | 12.67 ± 1.24 (94.1; ns) | 13.32 ± 0.91 (99.0; ns) |
| Keto acid | amount of keto acid (pmol (mg FW)<sup>-1</sup>); mean ± SEM (% compared to WT; significance level) |  |  |
|  | WT | #3 | #4 |
| KIC | 0.25 ± 0.03 | 3.52 ± 0.12 (1396.7 ± 45.8; ***) | 3.84 ± 0.60 (1525.3 ± 238.2; ***) |
| KIV | 0.39 ± 0.05 | 2.10 ± 0.10 (534.7 ± 25.7; ***) | 2.88 ± 0.54 (731.5 ± 136.2; ***) |
| KMV | 0.14 ± 0.03 | 0.61 ± 0.06 (436.9 ± 39.6; ***) | 0.89 ± 0.12 (636.4 ± 82.7; ***) |

## Acknowledgements

We would like to thank Elke Ströher and Harvey Millar for providing the knock-down line *GRXS15^amiR^* as well as Nicolas Rouhier, Olivier Keech and Jim Whelan for providing antibodies. We thank Philippe Fuchs, Stefanie Müller-Schüssele and Nicolas Rouhier for helpful discussion and critical reading of the manuscript.

